# Tracking Sleep-Linked Brain Fluid Dynamics Using Modified fNIRS: A Novel Noninvasive Window into Glymphatic Function

**DOI:** 10.1101/2025.10.27.684357

**Authors:** S Gupta, F Amyot, W Coon, A Pollatou, L Skeiky, S Seenivasan, K Kenney, A Lee, M Lettieri, A Penafiel, S Yalewayker, E Cheraghpour, P Sheth, E Metzger, C Scholl, JK Werner

**Affiliations:** Uniformed Services University, Bethesda, Maryland; The Geneva Foundation, Tacoma, WA; Walter Reed National Military Medical Center, Bethesda, Maryland; Johns Hopkins Applied Physics Laboratory, Laurel, Maryland; University of Pittsburgh Swanson School of Engineering, Pittsburgh, PA

## Abstract

**Background:** The glymphatic system, a brain-wide perivascular and interstitial waste and signal transport pathway for cerebrospinal fluid (CSF) to exchange with interstitial fluid (ISF), has pronounced activity during non-rapid eye movement (NREM) sleep and has been implicated in the pathophysiology of traumatic brain injury, Alzheimer’s disease, and mood disorders. However, direct measurement in humans is limited because current imaging methods rely on intrathecal contrast-enhanced magnetic resonance imaging (MRI), which is unsuitable for routine or naturalistic sleep studies. The absence of real-time, noninvasive monitoring methods that allow for natural sleep poses a major barrier to advancing glymphatic research in clinical settings.

**Methods:** To address this barrier, we developed a non-invasive functional near-infrared spectroscopy (fNIRS) forehead array in a wearable headband using non-standard wavelengths to allow for better sensitivity for water measurement. We monitored cortical blood and water dynamics during overnight sleep, quantifying oscillatory patterns of oxyhemoglobin (HbO) and water across sleep stages within the outer layers of the frontal cortex, subarachnoid space, and scalp. A component of the extracted water metrics is hypothesized to serve as a proxy for glymphatic transport without the need for contrast agents or surgical intervention.

**Results:** Our results demonstrate water concentrations were highest in SWS (d = 1.93, *p*=0.0002), while HbO concentrations also showed a modest elevation (d = 1.06, ns). Additionally, low-frequency oscillations (LFOs) of both water and HbO signals exhibited distinct dynamics of suprathreshold envelope peak (SEP) frequencies during NREM stages (N2 and N3) as compared to REM and wake, with effect size (d= 1.31, *p* = 0.003) for water. Interestingly, these water-derived metrics correlate with EEG slow-wave activity, linking fluid-sensitive oscillations to established electrophysiological markers of sleep depth. These findings indicate that water-sensitive oscillatory processes are selectively amplified during deep sleep and scale with EEG-defined sleep depth, consistent with a role for glymphatic-related fluid transport in human sleep.

**Conclusion:** We report novel cortical water shift parameters that are robustly sensitive to sleep stage transitions via a wearable, non-invasive, scalable headband, consistent with predicted glymphatic activity. Future work will cross-validate this method with MRI and other techniques.

**Disclaimer:** The views, information or content, and conclusions presented do not represent the official position or policy of, nor should any official endorsement be inferred on the part of, the Uniformed Services University, the Department of War, the U.S. Government, or Walter Reed National Military Medical Center.

## 1. Introduction

### 1.1. Background: The Glymphatic System and Brain Clearance

The glymphatic system is hypothesized to play a critical role in maintaining brain homeostasis, transporting signaling molecules and metabolic waste by facilitating the mixing of cerebrospinal fluid (CSF) with interstitial fluid (ISF) along a bulk flow gradient. This clearance mechanism is thought to be essential for removing neurotoxic substances such as amyloid-beta, alpha-synuclein, and tau, which accumulate during wakefulness and are strongly associated with the development of several neurodegenerative diseases (Hablitz & Nedergaard, 2021; Hauglund et al., 2020; Iliff et al., 2012). Non-rapid eye movement sleep (NREM) amplifies this fluid exchange, enabling more efficient transfer of signaling molecules and the clearance of harmful solutes from the brain’s interstitial fluid space (Van Hattem et al., 2025; Xie et al., 2013). Specifically, slow wave activity (SWA) has been strongly associated with glymphatic activity in both animal and human studies. For example, pioneering work by Xie and colleagues demonstrated that glymphatic flow increased by 60% during sleep vs. wakefulness, which correlated with large-amplitude, low-frequency electroencephalography (EEG) activity (SWA). In this work, animals were treated with anesthetics (ketamine/xylazine), which boosted SWA that was hypothesized to drive the expansion of interstitial space (∼60%) during sleep SWA, promoting the convective fluid osmotic flux by reducing neuronal firing and promoting vascular pulsatility - a putative driver of CSF-ISF exchange (Xie et al., 2013). In humans, emerging evidence suggests that SWA-rich NREM correlates with larger CSF oscillations in the fourth ventricle - hypothesized to relate to improved glymphatic dynamics (Fultz et al., 2019). Overall, SWA is considered a key physiological driver of glymphatic activity across species. Despite this strong evidence, noninvasive methods to directly monitor SWA-linked water flux in the human brain during sleep are lacking, underscoring the need for new measurement approaches.

### 1.2. Cerebral Blood Flow and Cerebral Blood Volume and Their Role in Sleep and CSF Flow

A key physiological driver of glymphatic clearance is cerebral blood flow (CBF), the volume of blood passing through the brain per unit of time, a direct function of cerebral blood volume (CBV). The brain’s high metabolic demand necessitates tight regulation of CBF and CBV via neurovascular coupling. This tight temporal linkage between neural activity and vascular physiology ensures the delivery of oxygen and nutrients to neurons and glia (Phillips et al., 2016). Arterial and venous pulsatility increases the hydrostatic pressure gradient in the vascular compartment, leading to interstitial water flux. Vasodilation or arousal can alter vascular compliance and increase CBV to amplify the perivascular pumping mechanics (Bohr et al., 2022). Neuronal activation, postural changes, smooth muscle contractions, or tissue compression also contribute to the pressure gradients, driving CSF into the perivascular space and facilitating its exchange with ISF (Fultz et al., 2019; Phillips et al., 2016; Rasmussen et al., 2022). These vascular pulsations occur across multiple frequency bands that have recently been linked to glymphatic function. Very-low-frequency vascular oscillations (VLFOs), approximately 0.01-0.06 Hz, are thought to be the drivers of B-waves. CSF pulsations originally observed by Lundberg, proposed to be the same oscillations driving glymphatic activity (Newell et al., 2022). Recently, in rodents, VLFOs were shown to be driven by the locus coeruleus, correlating with glymphatic exchange (Hauglund et al., 2025a; Lundberg, 1960). Low-frequency oscillations (LFOs, or Mayer waves), approximately 0.05-0.2 Hz, are driven by sympathetic activity and are significantly modulated during sleep and are thought to also contribute to CSF flow.

Supporting this mechanistic link, Yang and colleagues demonstrated that low-frequency vascular oscillations (VLFOs and LFOs together) during wakefulness drive bidirectional CSF flow in the fourth ventricle via CBF shifts (Yang et al., 2022). Extending this work, Nair et. al showed that systemic vasomotor LFOs continue to drive CSF flow across both wakefulness and NREM sleep, while SWA contributes an additional sleep-specific modulation (Nair et al., 2023). Thus, neural oscillations in the locus coeruleus and sympathetic circuitry are hypothesized to drive CBV and CBF fluctuations, the motive force for glymphatic activity. Hence, both CBF and CBV are intimately linked to water flux in the brain via their effects on vascular pulsatility, pressure gradients, and perivascular fluid dynamics. These interactions are likely critical for maintaining brain homeostasis and play a central role in waste clearance through the glymphatic system, especially during sleep (Fatima et al., 2025; Li et al., 2024). However, CBV and CBF detection methods focus on indirect hemodynamic measurement without the ability to isolate CSF-ISF flux, highlighting the need for optical tools that can distinguish blood from water and the mechanism between these vascular pulsatility and glymphatic clearance

### 1.3. The Gap: Lack of noninvasive tools to measure water-based fluid dynamics associated with glymphatic function during natural sleep

While contrast-enhanced magnetic resonance imaging (MRI) studies have confirmed sleep-associated increases in glymphatic activity (Eide & Ringstad, 2015, 2021; Taoka & Naganawa, 2020), they rely on intrathecal tracers and are unsuitable for longitudinal or naturalistic sleep research due to poor temporal resolution of the hemodynamic response in BOLD signals and participant discomfort. Notably, functional magnetic resonance imaging (fMRI) studies provide indirect evidence of glymphatic involvement by capturing CSF oscillations in the fourth ventricle that are tightly coupled with NREM sleep (Fultz et al., 2019; Helakari et al., 2022; Picchioni et al., 2022). These oscillations are hypothesized to drive convective fluid exchange by synchronizing arterial pulsations, CSF inflow, and interstitial clearance (Ladrón-de-Guevara et al., 2022; Lewis, 2021). However, such measurements are constrained by the uncomfortable conditions of the physically constrained MRI environment, making it challenging to study natural physiological sleep. Even when used, this approach remains too invasive for routine application and lacks the temporal resolution to capture the mechanism of dynamic, sleep-linked fluid exchange, leaving a significant translational gap in understanding human glymphatic function. Interestingly, in 2019 Meghdadi and colleagues developed a portable method to track electrical impedance during sleep, recapitulating older findings of impedance fluctuations in NREM sleep using a single carrier frequency (Meghdadi et al., 2019). In 2025, Dagum and colleagues used a more advanced technique, electrical impedance spectroscopy (EIS) with two in-ear electrodes, showing that reductions in brain parenchymal resistance change with sleep stages and correlate with MRI IV contrast migration into the cortex - a potential marker for both changes in CBF and glymphatic activity (Dagum et al., 2025a). Using a wearable, wireless device, Dagum and colleagues linked lower resistance to increased fluid content in the brain parenchyma and glymphatic exchange, mirroring prior findings in animal models. Importantly, EEG delta power, reduced beta power, and lower heart rate were associated with improved glymphatic clearance. Importantly, the impedance approach is unable to distinguish water changes compared to blood - nor can it localize the effects it is measuring; rather, they are global changes in impedance, nonspecific to water, which may explain its modest signal-to-noise ratio and effect size. Impedance changes cannot observe extracellular water (ISF/CSF) separately from blood, potentially limiting the dynamic range of the impedance measurement (Dagum et al., 2025a). To measure glymphatic physiology in higher resolution and greater specificity, other techniques are needed.

### 1.4. The Promise of Optical Imaging for the Unmet Need

Functional near-infrared spectroscopy (fNIRS) offers a promising non-invasive approach to study sleep-dependent fluid dynamics, which includes glymphatic activity. Unlike fMRI, fNIRS is capable of using photon absorbance spectroscopy to estimate relative concentrations of specific molecules, classically resolving oxyhemoglobin (HbO) and deoxyhemoglobin (Hb), among other molecules. Fantini and colleagues demonstrated that spontaneous low-frequency oscillations (LFOs) in HbO/Hb are attenuated during NREM sleep and rebound during REM and wake, reflecting stage-dependent modulation of cerebral blood volume, while phase differences between HbO and Hb oscillations served as surrogates for changes in blood flow (Pierro et al., 2012). Oniz and colleagues further validated fNIRS sensitivity to sleep-related cerebrovascular regulation by observing distinct HbO/Hb concentration fluctuations during NREM–REM transitions (Oniz et al., 2019). Extending beyond hemodynamics, Myllylä and colleagues applied longer, water-sensitive wavelengths to fNIRS, providing the first evidence that cortical water dynamics can be tracked noninvasively in awake humans (Myllylä et al., 2018). Most recently, Yoon and colleagues (Yoon et al., 2025) demonstrated that fNIRS can capture gross (NREM vs REM) sleep-state-dependent changes in frontal cortex water content.

As shown previously, we also expect fNIRS-measured Δwater to increase with deeper sleep. Some other key differences in our approach include: 1) the use of an array of optodes and detectors across the forehead to assess laterality; 2) a NIRS analysis pipeline that measures and reports changes in both blood (HbO) and water through the night 3) evaluation of HbO/Hb concentration differences across individual sleep stages; 4) analysis of the EEG slow wave activity, the hemodynamic response, and water response; 5) a venous blood sampling just before and after sleep to measure biomarkers associated with glymphatic activity (in process). In this study, we test a real-time, noninvasive tracking of sleep-dependent water and blood flux using a scalable fNIRS wearable headband.

## 2. Methods

### 2.1. Participant recruitment and selection

Nine healthy volunteers were recruited from clinics at Walter Reed National Military Medical Center (WRNMMC) as part of a larger parent study focused on sleep physiology. Eligible participants were 18-55-year-old Department of Defense (DoD) beneficiaries with sleep-related complaints necessitating overnight polysomnography (PSG).. Exclusion criteria included severe insomnia, habitual insufficient sleep, prior stroke or transient ischemic attack (TIA), uncontrolled migraine, congestive heart failure, pulmonary hypertension, kidney disease, caffeine intake >400 mg/day, Body Mass Index (BMI) >34, clinically unstable psychiatric conditions, pregnancy, moderate or severe obstructive sleep apnea (apnea–hypopnea index (AHI) >15), and any history of Traumatic Brain Injury (TBI). This work was conducted under a protocol approved by the Uniformed Services University Institutional Review Board.

### 2.2. Clinical Overnight PSG and fNIRS data acquisition

Participants underwent overnight PSG in an American Academy of Sleep Medicine (AASM) accredited sleep laboratory, where physiological signals were continuously monitored, including EEG, electrooculogram (EOG), electromyogram (EMG), electrocardiogram (ECG), respiratory effort, airflow, oxygen saturation (SpO_2_), and limb movements. The Nihon-Kohden Polysmith 13 was used for PSG collection, including EEG tracings for each participant at a 500Hz sampling frequency. Sleep stages were determined both manually by registered PSG technologists and automatically, using Yet Another Spindle Algorithm (YASA) (Vallat & Walker, 2021). Since delta power calculations were also used with the YASA algorithm, and because agreement with certified technologists was >90%, we elected to use YASA for sleep stage scoring to correlate delta power of slow waves with fNIRS.

In parallel with PSG, we employed a custom-engineered continuous-wave fNIRS system consisting of light-emitting diode (LED) sources and photodiode detectors to record throughout the night. The fNIRS headband was positioned over the prefrontal cortex, ensuring proper sensor placement and minimal interference with PSG electrodes. Dual-wavelength near-infrared light sources (860 and 980 nm) were used to monitor changes in absorbed light intensity continuously. The system operates at a sampling rate of 10 Hz for each source, multiplexing to ensure no overlap of photons. The optical sources had an approximate output power of 210 mW/cm^2^, well within the safety limits for tissue exposure. Photodiode detectors were located 3 cm, 4 cm, and 5 cm apart from the sources, consistent with standard source-detector distances demonstrated to create photon paths that pass through the outer 0.1 to 0.5 cm of the cortex (Delpy et al., 1988; Durduran et al., 2010; Strangman et al., 2013a). Signal quality was assessed in real-time, and measurements were continuously monitored by trained research personnel to minimize motion artifacts and maintain optimal data quality. The highest signal-to-noise (SNR) was observed in the channels with a 3cm source-detector (S/D) separation. These 8 channels yielded a median SNR of 19dB (inter-quartile range: 10dB to 28dB). Because of the greater signal quality, we used the data from the 3cm S/D channels exclusively for our physiological analysis. The full SNR profile across all S/D channel lengths is provided in Supplementary Figure 1.

All equipment had been configured to optimize participant comfort and safety during overnight recording. Time synchronization between the PSG and fNIRS systems was performed at the start of each session by utilizing a DC output trigger box to enable precise alignment of electrophysiological and hemodynamic data for subsequent analysis. Clinical staff remained on site and available throughout the night to address any technical issues or participant needs, ensuring adherence to safety protocols and high-fidelity data acquisition.

### 2.3. Preprocessing of Near-Infrared Spectroscopy (NIRS) Data

fNIRS data underwent a comprehensive preprocessing pipeline to ensure signal quality and minimize artifacts, which included motion correction, detrending, and filtering to isolate the signal output(Figure 2). Motion artifacts were identified using the moving standard deviation (MSD) (Scholkmann et al., 2010), computed for each time point across the signals. A sliding window approach (299s window) was applied to calculate MSD, and segments exceeding predefined thresholds were flagged and masked. This ensured that noisy regions caused by motion were excluded from further analysis.

**Figure 1.**
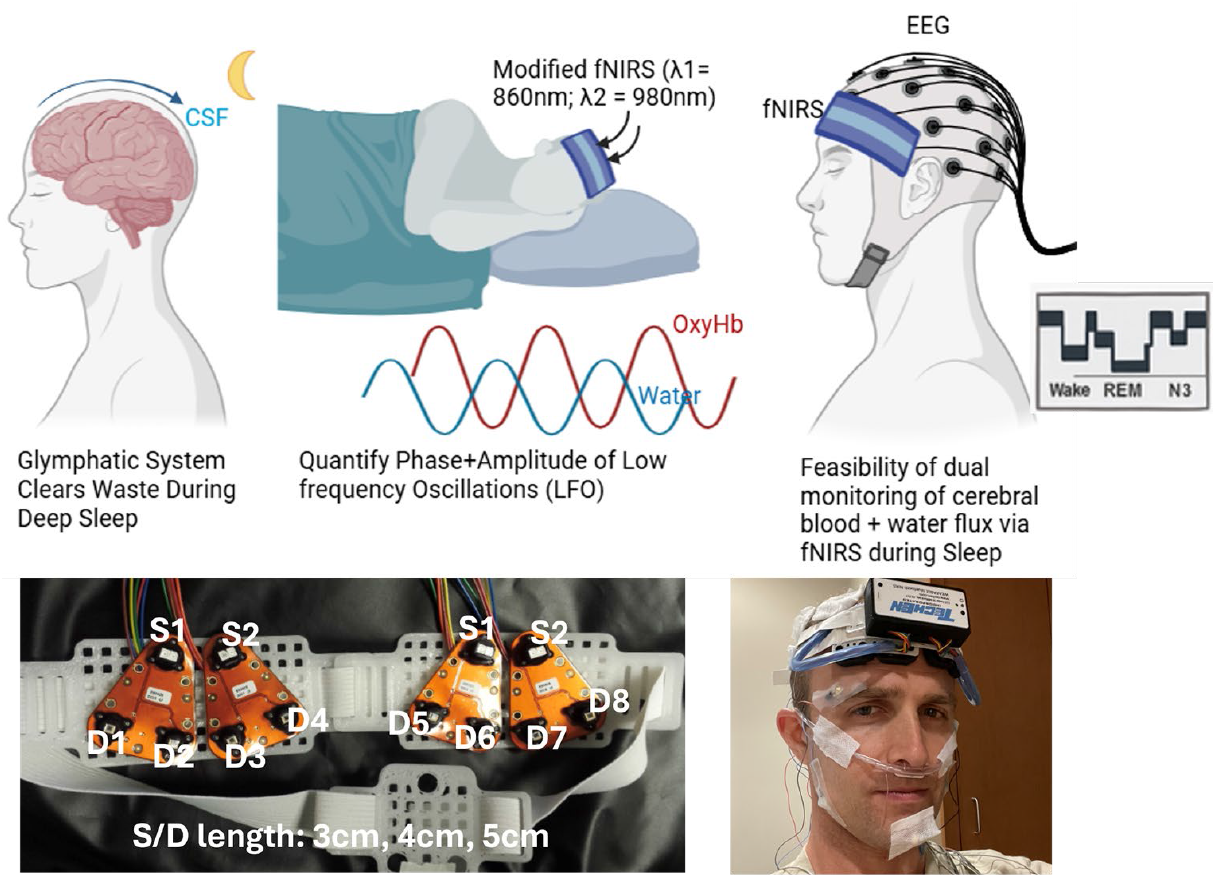
Conceptual overview and experimental setup for dual monitoring of cerebral blood and water dynamics during natural sleep using modified fNIRS. (Top row) A modified functional near-infrared spectroscopy (fNIRS) device with dual wavelengths (λ_1_ = 860 nm for oxyhemoglobin, λ_2_ = 980 nm for water absorption) is used during overnight polysomnography (PSG) to quantify low-frequency oscillations (LFOs) in both hemoglobin and water signals. The phase and amplitude of these oscillations are extracted to assess cerebral hemodynamics and water flux across sleep stages. (Bottom row) Photos of the custom fNIRS sensor array and photo of PI (corresponding author) showing setup demonstrating the feasibility of simultaneous EEG and fNIRS recordings during sleep

**Figure 2:**
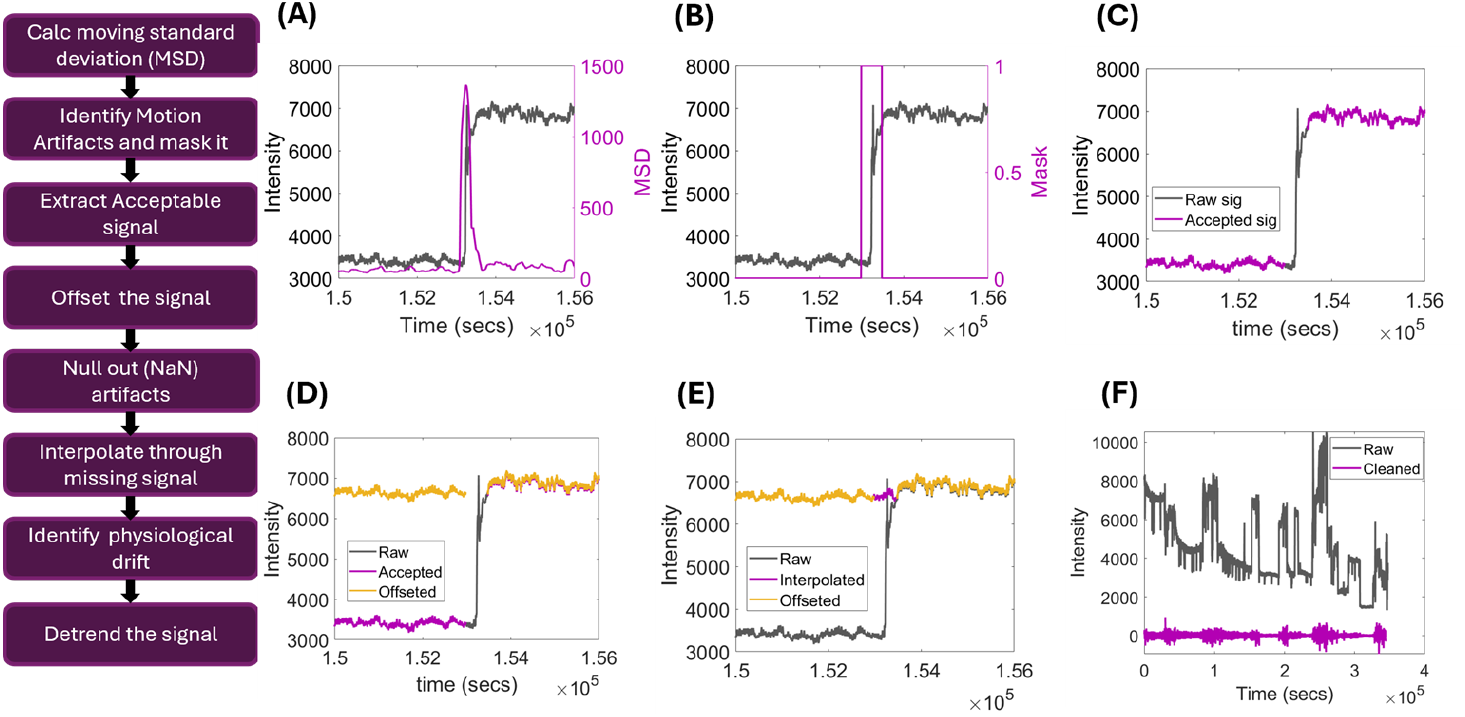
Preprocessing pipeline for fNIRS signal denoising and artifact correction. (Left) Schematic flowchart outlining the sequential steps of the preprocessing pipeline. (Right) Panels A–F illustrate the progression of signal cleaning; (A) Raw intensity trace with MSD (purple) showing a spike due to motion artifact; (B) Masking of high-MSD region identified as artifact; (C) Extraction of artifact-free signal segment; (D) Offset correction applied to accepted segment; (E) Interpolation across missing segments using Fourier-based estimation; (F) Final cleaned and detrended signal, preserving physiological oscillations while removing artifacts and low-frequency drift. This preprocessing pipeline enhances signal quality for downstream analysis of cerebral hemodynamics and water flux in sleep fNIRS data.

After masking, acceptable signal segments were extracted for preprocessing. Baseline shifts were corrected using an offset adjustment, normalizing the signal across channels and subjects for comparability. Artifact-affected segments, including motion-induced noise and missing data, were replaced with NaN values to prevent their influence on subsequent calculations. Missing segments were interpolated using Fourier interpolation, ensuring smooth signal recovery while preserving physiological patterns.

Long-term drifts and very low-frequency fluctuations (<1mHz), presumed to reflect instrumental drift, were removed by fitting and subtracting a first-order polynomial from the raw signal. The detrended data were subsequently band-pass filtered within the frequency range of interest, thereby minimizing residual noise and isolating physiologically relevant activity.

The preprocessing pipeline was implemented in MATLAB 2024b (MathWorks Inc.) using custom scripts and built-in signal processing functions. It was validated on a subset of the data for robustness and reproducibility. Visual inspections were conducted at multiple stages, and signal amplitude thresholds were applied, with values exceeding 500 (a.u.) considered indicative of high-quality signals. Approximately 30–40% of the data were rejected on average due to noise and motion artifacts. This pipeline enabled the extraction of high-quality, artifact-free NIRS signals, facilitating accurate analyses of cerebral hemodynamics and physiological processes.

### 2.4. Estimation of Oxygenated Hemoglobin (HbO) and Water Concentrations Using the Modified Beer-Lambert Law (MBLL)

The preprocessed NIRS data were used to estimate HbO and water concentrations based on the MBLL, which quantifies chromophore concentrations by analyzing wavelength-dependent light attenuation through tissue. Following the previously described cleaning pipeline, the NIRS data consisted of light intensity measurements at 860 nm and 980 nm, selected for their sensitivity to HbO and water content because they flank the isobestic point.

The MBLL is expressed as:

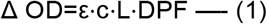

Where Δ OD = Change in optical density (attenuation); ε = Extinction coefficient of the chromophore at a given wavelength; c = Chromophore concentration; L = Optical path length (S/D lengths of 3cm, 4cm, and 5cm); DPF = Differential Pathlength Factor (accounting for tissue scattering). Using equation (1), changes in optical density (ΔA) were converted into concentrations of HbO and water by solving for c. Extinction coefficients (HbO at 860nm: 1092 M^-1^cm^-1^ and at 980nm: 1128 M^-1^cm^-1^; H2O at 860nm: 0.043 M^-1^cm^-1^ and at 980nm: 0.43 M^-1^cm^-1^) for these chromophores were taken from published literature (HOMER) as compiled by Scott Prahl (Prahl, n.d.), referring to the work from (Kollias & Gratzer, 1999). For simplicity, the DPF was held constant at 1 across wavelengths and participants, consistent with relative change analysis.

### 2.5. Spectral Decomposition of HbO and H_2_O oscillations

After converting raw intensities to concentrations with MBLL, we extracted sleep-stage–specific oscillations of HbO and H_2_O and quantified their spectra. Hypnograms were upsampled from 30-s epochs to 10 Hz to match the fNIRS data sampling frequency. For each stage (N2, N3, REM, Wake), the corresponding samples from the concentration time series were indexed. We analyzed the 3cm S/D separation channel for spectral decomposition. Using Chronux ‘*mtspectrumc*’, we computed the power spectral density. The raw spectrum was lightly smoothed by a Savitzky–Golay filter to emphasize broad peaks without inflating bandpower. Spectra were restricted to bandpass filtered from 0.001–1.2 Hz and normalized to unit area, giving a probability-like spectrum per stage. Bandpower (area-under-curve) was then computed in pre-registered bands (LFO: 0.005–0.2 Hz, RFO: 0.2–0.4 Hz, CFO: 0.6–1.2 Hz). The same per-stage spectra were exported to Python and passed to the FOOOF algorithm to model the broadband 1/f background and the residual oscillatory “peaks.” FOOOF yields the aperiodic component (background) and the periodic (peak) component; the latter is plotted and shown in results as the dashed “Mean Periodic Component” in Figure 3. Group means (±SD across participants) are shown as shaded fills; the dashed line visualizes oscillatory energy after removing the 1/f background.

**Figure 3.**
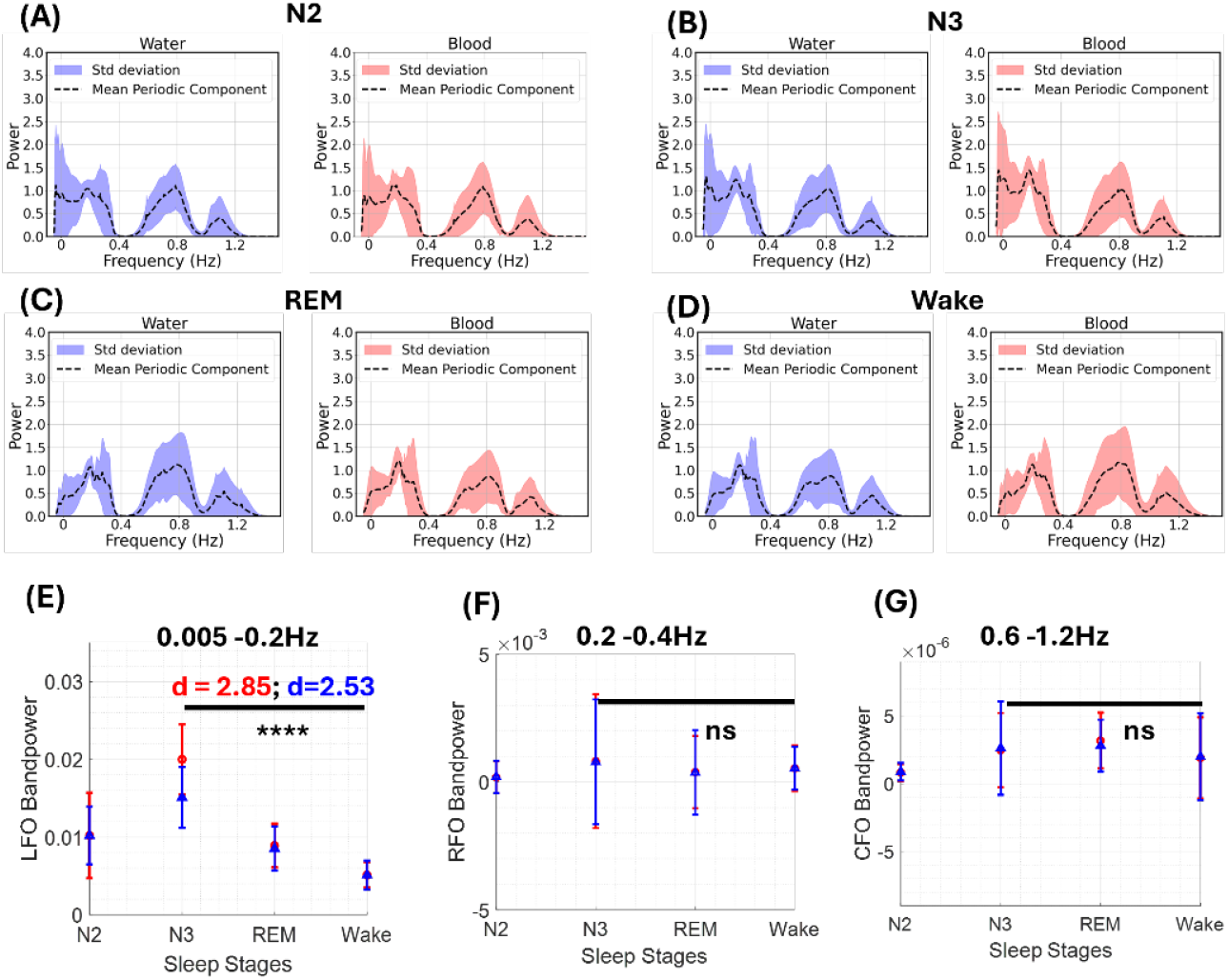
Power spectral density of water and oxyhemoglobin signals across sleep stages. (A–D) Frequency-domain analysis of cerebral water (left) and oxyhemoglobin (right) signals during N2, N3, REM, and Wake stages. Shaded regions represent standard deviation across subjects, and dashed lines represent the mean periodic component within each frequency bin. Prominent spectral peaks were observed in the low-frequency range (0.005–0.2 Hz), with additional components in respiratory (0.2–0.4 Hz) and cardiac (1.0– 1.2 Hz) frequency ranges. (E) Quantified bandpower in the low-frequency range showed significantly greater power during N3 sleep in both water (blue) and HbO (red) signals compared to other stages (Cohen’s *d* = 2.85 for water, *d* = 2.53 for blood; \*\*\*\**p* < 0.0001). (F–G) No significant differences were observed across stages in the respiratory frequency (RFO: 0.2–0.4 Hz) or cardiac frequency (CFO: 1.0–1.2 Hz) bands.

### 2.6. fNIRS metrics Calculation from MBLL derived concentrations of HbO and H2O

We converted raw fNIRS intensity to concentration changes using a two-wavelength MBLL workflow. For each file, channels 1–16 (≈860 nm) and 17–32 (≈980 nm) were offset (+10 000) and transformed to optical density, then solved for Δ[HbO] and Δ[H_2_O] with extinction coefficients supplied to the MBLL custom function written in MATLAB. Channels with ≥50% missingness at either wavelength were excluded. We then characterized fast dynamics via the analytic signal. For each usable channel, we computed the Hilbert transform to obtain the instantaneous phase (unwrapped) and amplitude, normalizing amplitudes per channel. To compare blood and water dynamics, we focused on 3 cm S/D and 4 cm S/D separation channels and formed sample-wise HbO– H_2_O phase differences using the circ_dist function from the circular statistics toolbox available in MATLAB (Berens, 2009), followed by a circular mean across the selected channels (circ_mean) with NaN masking. In parallel, we summarized amplitude by taking the per-sample median across channels for HbO and H_2_O. Finally, we aligned these signals to sleep architecture. Hypnograms were standardized, edge-trimmed, and upsampled by replication (factor = 300) to match signal length. Stage indices (N1, N2, N3, REM, Wake) were used to extract stage-specific segments. For each stage, we computed circular-mean phase difference (degrees), median amplitudes for H_2_O and HbO, and peak metrics using the built-in ‘findpeaks’ function in MATLAB with adaptive prominence (mean + 1×SD) reporting median peak height and normalized peak counts.

In our peak analysis, we first derived a clean amplitude envelope from each HbO/H_2_O concentration trace by taking the analytic signal via the Hilbert transform and using its magnitude A(t)=∣x(t)+jH{x(t)}∣; envelopes were masked where channels are invalid and, for robustness, pooled across the selected 3cm S/D-separation channels using the sample-wise median. For each sleep stage, we then detected suprathreshold envelope peaks with MATLAB’s ‘findpeaks’ function, using a stage-adaptive threshold T = μ+ kσ computed on the stage-restricted envelope (where μ = mean, σ = SD, and k = 1). This was applied as the MinPeakProminence argument in the findpeaks function (how much a peak stands out locally). To avoid double-counting, we used a physiological minimum peak distance (1s in samples). From the detected peaks, we reported peak counts as the suprathreshold envelope peak (SEP) rate. All computations are performed per stage after masking/trim steps, so thresholds and counts are adapted to the distribution of the envelope within each stage rather than using a global cutoff.

### 2.7 Water Signal Filtering

To reduce crosstalk between the water signal and hemoglobin-related changes, we adopted the linear minimum mean square estimator (LMMSE) filtering approach described by Yoon and colleagues (Yoon et al., 2025). This static filter method isolates the water component independent of HbO by removing the plasma-related fraction that covaries with hemoglobin concentration. Following their framework, the measured water content signal (ΔH_2_O measured) was decomposed into two components: (i) the plasma-related portion proportional to changes in hemoglobin concentration (ΔC of HbO) and (ii) the residual component representing brain water dynamics unrelated to HbO. The filtered water signal was therefore obtained as:

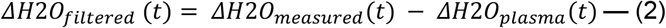

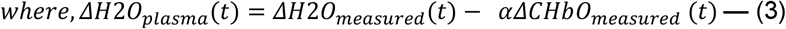

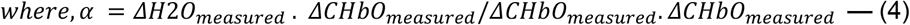

### 2.8. EEG Data Processing for Spectral Power Analysis

The raw EEG tracings were imported from EDF files using MNE-Python. If the Nyquist frequency was above 60Hz, the EEG recordings were then notch filtered at 60Hz using Python SciPy’s ‘iirnotch’ at a quality factor of 30, defining a narrow bandwidth of 2 Hz (Dagum et al., 2025). Continuous data were then bandpass filtered between 0.3 Hz and 49.5 Hz using a finite impulse response filter with a Hann window (‘firwin’). Filter lengths were set at 2401 taps for the high-pass edge and 501 taps for the low-pass edge, following the standard approximation N=4/ΔF, where ΔF= transition band / nyquist frequency (Dagum et al., 2025). This design ensured sharp spectral transitions and preserved high-resolution signal components by implementing a high-order digital filter with a narrow and precise stopband.. The filter was applied bidirectionally using SciPy’s ‘filtfilt’ to achieve zero phase distortion. Data were resampled to 100Hz using MNE ‘resample’ to reduce computational load and storage. Finally, preprocessed EEG data were stored in the MNE native format (.FIF file, functional imaging file) to enable efficient indexing and loading for subsequent time-frequency analyses.

Preprocessed FIF files were loaded using MNE-Python, and all EEG channels were converted to microvolts. The data was segmented into 5-second windows using the automated YASA ‘sliding_window’ function, resulting in a 3D array of shape ‘epochs, channels, samples**’** (Vallat & Walker, 2021). At the resampled frequency of 100Hz, each epoch contained 500 samples. The hypnogram was repeated at each stage to match the 5-second epochs and then padded with the last stage to match the epoch length. The C4-M1 channel was selected for analysis, and it underwent two layers of artifact rejection. The artifact rejection methodology followed the algorithm utilized in the paper. First, a peak-to-peak amplitude criterion was applied, rejecting epochs with voltages exceeding ±350 μV. Second, the power spectral density (PSD) of each epoch was calculated using Welch’s method, SciPy’s ‘signal.welch’, using a 10-second sliding window and 50% overlap. Epochs exceeding a maximum PSD of 1000 µV^2^/Hz were removed to minimize the influence of high-frequency noise and non-physiological drifts. The relative bandpowers were derived from the cleaned C4-M1 channel PSD using YASA’s ‘bandpower_from_psd_ndarray’ function. Frequency bands of interest were defined as Delta (0.5-4Hz), Theta (4-8 Hz), Alpha (8-12 Hz), Sigma (12-16Hz), and Beta (16-30 Hz). Bandpower estimates were normalized to the total power across these ranges using Simpson’s integration (scipy.integrate.simps). Average relative bandpowers were then computed for each sleep stage.

## 3. Results

### 3.1 Participant Demographics and PSG Data

In Table 1, we summarize the demographic and sleep characteristics of the study sample (N = 9 participants, 13 total recordings). Participants had a median age of 42 years (IQR = 11), with most identifying as White (66.7%) and male (77.8%). The median AHI was 3, indicating a low sleep-disordered breathing burden. Habitual sleep timing derived from actigraphy showed a median midsleep point at 03:08am and a habitual sleep duration of 7.8 hours. PSG metrics indicated a median total sleep time (TST) of 351 minutes, with stage distributions of NREM1 (7.57%), NREM2 (56.02%), NREM3 (17.82%), and REM sleep (19.16%). Table 1 displays demographics of the cohort; participants were primarily middle-aged White males (77.8%) with generally healthy sleep architecture and low AHI, providing a stable baseline for analyzing physiological signals across sleep stages.

**Table 1.** Demographics and PSG results of study participants. Values are presented as median (Mdn) ± interquartile range (IQR) unless otherwise indicated. A total of N = 9 study participants were included in this analysis; however, n = 13 records were analyzed as four participants contributed two records of usable data each. For each participant, age, sex, race, apnea–hypopnea index (AHI) is reported alongside their habitual midsleep point and sleep duration derived from Oura Ring, and polysomnography (PSG) metrics, including total sleep time (TST) and non-rapid eye movement (NREM) stages 1–3, as well as rapid eye movement (REM) sleep.

| Study Participants |  | Demographics |  |  | PSG Characteristics |  |  |  |  |  |  |  |
| --- | --- | --- | --- | --- | --- | --- | --- | --- | --- | --- | --- | --- |
| Participant<br>t<br>(n = 9) | #Records<br>(n=13) | Race<br>(%) | Age<br>Mdn(IQR) | Sex<br>(%) | Habitual<br>Midsleep<br>Point hh:mm<br>(24 h)<br>Mdn(IQR) | Habitual<br>Sleep<br>Duration<br>(min)<br>Mdn(IQR) | AHI<br>Mdn(IQR) | TST<br>(min) | NREM<br>Stage 1<br>(min) | NREM<br>Stage 2<br>(min) | NREM<br>Stage 3<br>(min) | REM<br>(min) |
| **A**<br>Record 1<br>Record 2 | 2 | Black | 42 | M | 02:53 (72.4) | 483.8 | 10<br>2 | 231.5<br>284.0 | 23.5<br>21.5 | 124.5<br>179.5 | 38.0<br>43.5 | 45.5<br>39.5 |
| **B**<br>Record 1<br>Record 2 | 2 | White | 30 | F | 02:10 (97.5) | 502.2 | 2<br>2 | 347.0<br>375.5 | 13.5<br>22.5 | 195.5<br>195.5 | 71.5<br>96.0 | 66.5<br>61.5 |
| **C**<br>Record 1<br>Record 2 | 2 | White | 35 | M | 03:08 (70.6) | 473.4 | 12<br>12 | 276.0<br>394.5 | 16.5<br>20.5 | 173.0<br>243.0 | 49.5<br>63.5 | 37.0<br>67.5 |
| **D**<br>Record 1<br>Record 2 | 1 | N/A | 43 | F | 04:52 (70.7) | 464.6 | 1<br>2 | 498.0<br>495.0 | 39.0<br>26.5 | 314.5<br>282.5 | 41.0<br>74.0 | 103.5<br>112.0 |
| **E** | 1 | White | 44 | M | 03:50 (65.2) | 404.7 | 4 | 311.5 | 46.5 | 174.5 | 55.5 | 35.0 |
| **F** | 1 | White | 47 | M | 02:44 (112.1) | 471.7 | 7 | 360.0 | 18.5 | 161.0 | 65.5 | 115.0 |
| **G** | 1 | White | 25 | M | 05:10 (55.8) | 515.5 | 5 | 332.0 | 54.5 | 155.5 | 70.0 | 52.0 |
| **H** | 1 | White | 33 | M | 01:28 (82.0) | 466.4 | 0 | 394.0 | 36.0 | 182.5 | 92.5 | 83.0 |
| **I** | 1 | Mixed | 48 | M | 03:56 (138.2) | 442.3 | 3 | 351.0 | 32.0 | 179.0 | 49.5 | 90.5 |
| **All** | - | White: 6<br>(66.7%)<br>Black: 1<br>(11.1%)<br>Mixed: 1<br>(11.1%)<br>N/A: 1<br>(11.1%) | 42 (11) | Male: 7<br>(77.8%)<br>Female: 2<br>(22.2%) | 3:08 (72.4) | 471.7<br>(19.2) | 3 (5) | 351.0 (82.5) | 23.5 (15.5) | 179.5 (22.5) | 63.5<br>(22.0) | 66.5<br>(45.0) |

### 3.2. Frequency domain analysis of HbO and water signals

Filtered and cleaned HbO and water NIRS signals collected during PSG were sorted by sleep stage (excluding N1 due to the limited number of epochs) and transformed into the frequency domain to assess known physiologically relevant oscillatory bands and their changes specific to each sleep stage. As shown in Figure 3, peaks of spectral power were observed in the low-frequency oscillation band (LFO, 0.005–0.2 Hz), the respiratory frequency oscillation band (RFO, 0.2–0.4 Hz), and the cardiac frequency oscillation band (CFO, 0.6–1.2 Hz) which had a bimodal distribution. We quantified the bandpower of the signals (Figure 3, E-G) for each frequency band.

During NREM sleep (N2 and N3; Figure 3, A and B), both water and HbO signals exhibit a pronounced power in the low-frequency band (0.005–0.2 Hz) compared to REM and Wake (Figure 3, C and D).

Quantitative comparisons (Figure 3, E–G) reveal that bandpower in the low-frequency range is significantly higher during N3 than in all other stages, for both water and blood signals. In contrast, no significant stage-dependent changes were found in the respiratory or cardiac frequency bands.

### 3.3. Low Frequency Oscillation Analysis

We examined sleep water and HbO dynamics in greater detail in the LFO band. For this analysis, we computed the amplitude of the concentrations separately for each 30-second window, then averaged the results across the four sleep stages: N2, N3, REM, and WAKE. We initially included all water signals in our concentration analysis, finding mean concentrations are most elevated during N3, with a robust effect size (Supplemental Fig 2A \*\*\*\**p* < 0.0001, d = 2.38), and this signal trends positively with relative delta power but is not significant (Supplemental Figure 2B, r = 0.357, *p* = 0.23). We investigated whether regressing out the component of the water signal that oscillates in time with HbO - a method described recently by Yoon and colleagues (Yoon et al., 2025) - would enhance the relationship. Interestingly, this regression resulted in a robust improvement in the EEG delta power correlation with water concentration (Figure 4B, r = 0.63, *p* = 0.02). Between-stage concentration changes remained robust with this filtering approach, though the effect size was smaller as compared to our previous analysis without regressing out the water signal that oscillates with HbO (Figure 4A). Water mean concentration showed a non-significant association with the delta-to-beta ratio (Figure 4C). In contrast, HbO mean concentration shows no significant modulation across sleep stages (Figure 4D) and has no relationship with delta power or delta/beta ratio (Figure 4E-F).

**Figure 4.**
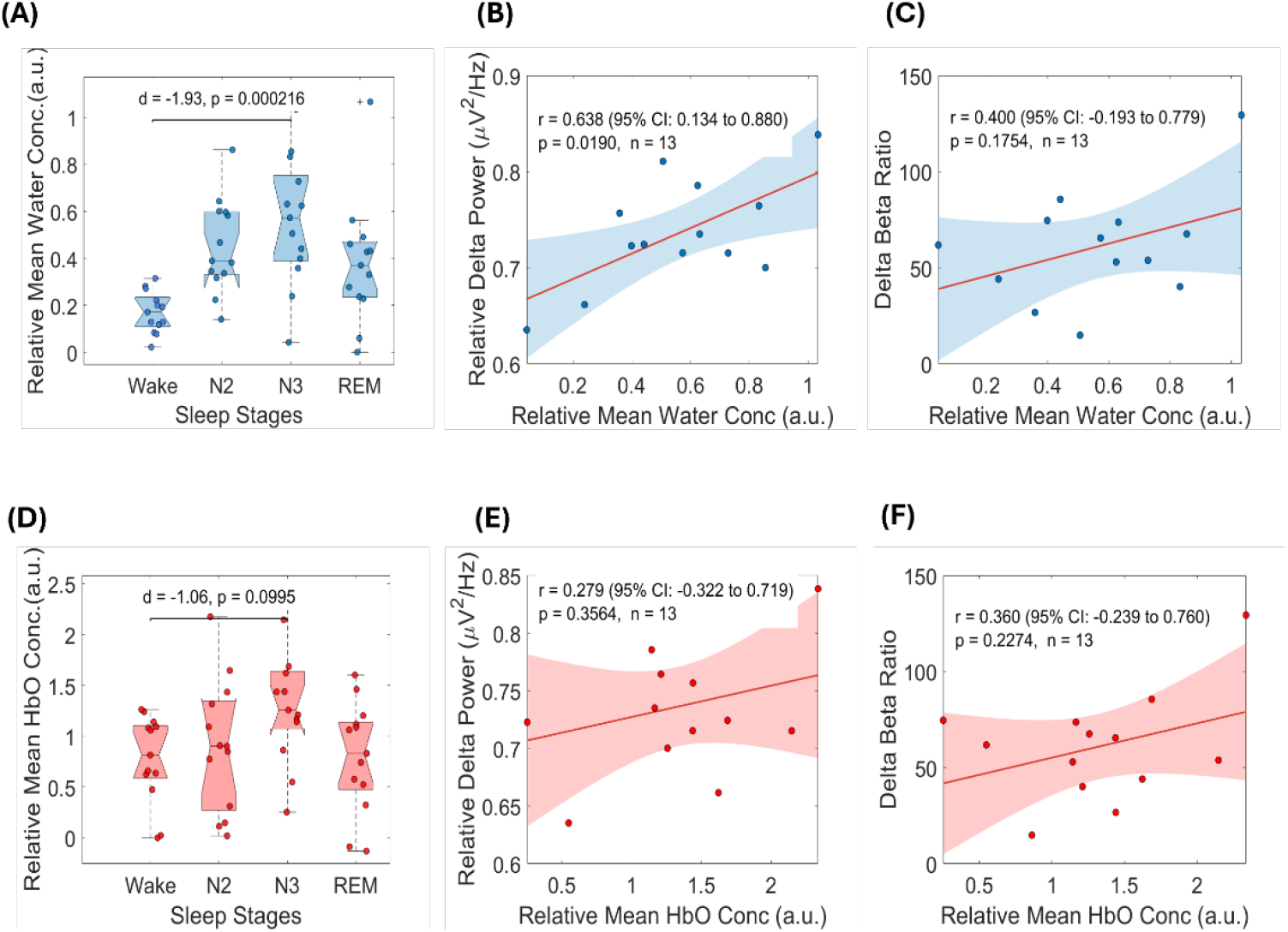
Water and HbO LFO dynamics compared across sleep stages and relationship to EEG power spectra in sleep and wake. (For all plots, N = 9 participants, n = 13 recordings). (A) H2O relative mean concentration is significantly elevated during N3 relative to other stages. (B) EEG relative delta power shows a strong positive trend with significant correlation with H2O mean concentrations. (F) Normalizing EEG delta with beta power demonstrates a slight positive association with H2O relative mean concentrations. (D) HbO mean concentration does not significantly differ across sleep stages or show a significant relationship with EEG delta or delta/beta power (D-F).

We also investigated if a focus on the larger water and blood concentration surges, defined by envelope peaks crossing a pre-defined threshold of *mean* ± 1 ∗ *standard deviation*, could better identify water flux. We measured suprathreshold water concentration envelope peak (SEP) rates across the sleep stages (Figure 5), observing that the H_2_O SEP rate was markedly higher during N3 compared to all other stages, with a very large effect size (d = 1.31, *p* = 0.003). By contrast, HbO SEP rate did not differ significantly across stages. When relating these measures to EEG spectral activity, H_2_O SEP rate during N3 showed a moderately positive association with relative delta power (r = 0.516, *p* = 0.07), though this trend did not reach statistical significance. Regressing out the HbO (vascular) related water signal did not significantly improve either the water or HbO relationship with EEG delta power or delta/beta power ratio. HbO SEP rate showed no meaningful relationship to delta power and demonstrated no significant association with the delta-to-beta ratio (r = -0.328, *p* = 0.274)

**Figure 5:**
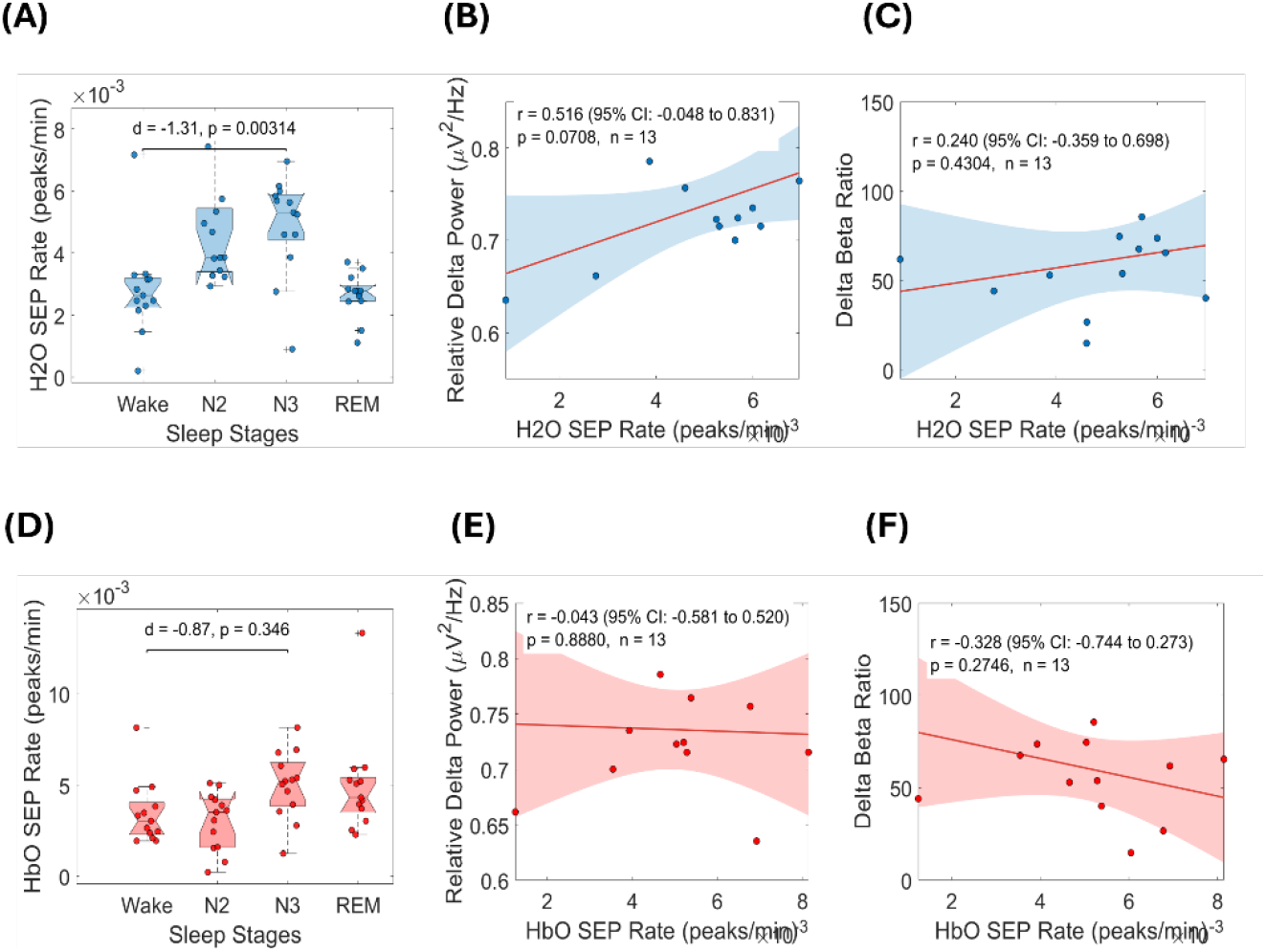
Peak oscillation dynamics of water and hemoglobin signals across sleep stages. (N = 9 participants, n = 13 recordings). (A) H_2_O suprathreshold envelope peak (SEP) rate is significantly elevated during N3 compared to other sleep stages. (B) H_2_O SEP rate at N3 shows a positive, near-significant correlation with relative delta power. (C) H_2_O SEP rate also trends positively with delta-to-beta ratio, although not significantly. (D) HbO SEP rate does not significantly differ across sleep stages. (E) HbO SEP rate shows no meaningful correlation with relative delta power.. (F) HbO SEP rate demonstrates a negative association with the delta-to-beta ratio, although not significant.

### 3.4. Respiratory and Cardiac Frequency Oscillation Analysis

We analyzed the data in the RFO and CFO frequency bands in addition to the LFO bands to see the nature of the signal in these frequencies of physiological oscillations. In panel A, H_2_O mean concentration is significantly larger during N3 sleep relative to other stages (d = 1.44, *p* < 0.01). HbO mean concentration changes are not statistically different across the stages. We then characterized the amplitude parameters in RFO, as we did for LFO, and examined the SEP rate of water and HbO oscillations across sleep stages, as shown in Supplementary Figure 4. While the water SEP rate was significantly higher in N3 relative to other stages (*p* < 0.05, d = 1.21), the HbO SEP rate was not significantly modulated by sleep stage. We then analyzed the data in the CFO band as shown in Figure 6 C-D. We see no stage-dependent modulation of concentration changes of water and HbO in the CFO band.

**Figure 6:**
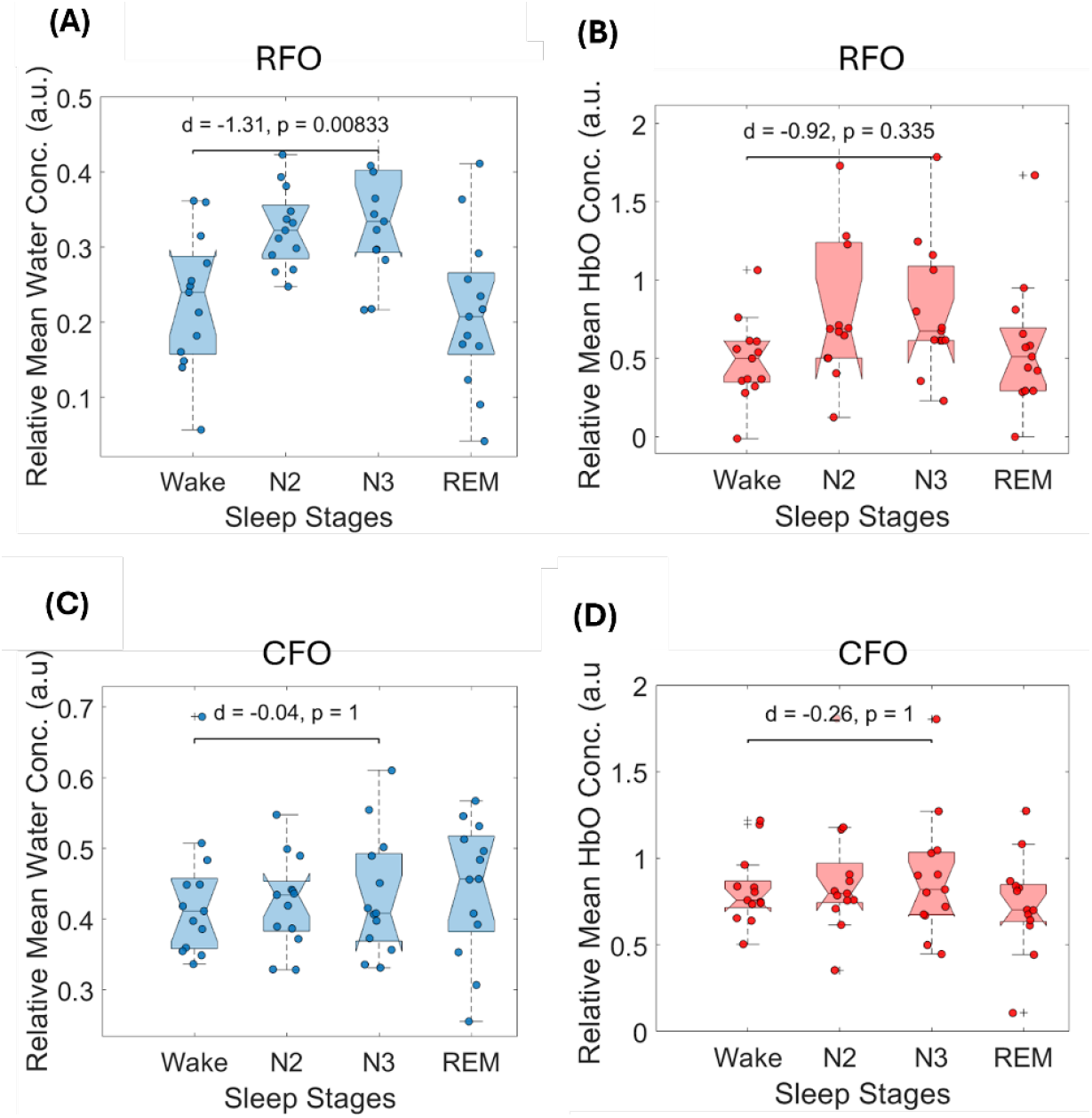
Mean concentration of H_2_O and HbO across sleep stages in the RFO band (0.2–0.4Hz; top row) and CFO band (0.6-1.2Hz; bottom row) (N=9 participants, n=13 recordings). (A) H_2_O mean amplitude shift was also elevated during N3 relative to Wake (*d* = 1.31, *p* = 0.008). (B) HbO mean amplitude shift showed a smaller effect (*d* = 0.92) and did not vary significantly across stages. (C-D) Amplitude dynamics in the cardiac-frequency range (0.6– 1.2 Hz) showed no significant differences across N2, N3, REM, or Wake. r

### 3.5. Night-to-night variability and signal symmetry

We computed the variability coefficients for multi-night participants while also comparing signal amplitude across the source-detector pairs on the left compared to those on the right for each participant. Intraclass correlation coefficients were low for most participants and are shown in Supplementary Figure 6. No significant differences were observed between left- and right-sided fNIRS measurements (Supplemental Figure 7).

## 4. Discussion

### 4.1. Temporal Decoupling of Cerebral Blood and Water Signals During NREM Sleep as a Marker of Cerebral Compliance and Glymphatic Function

We report a novel fNIRS approach to assess relative changes in brain compliance, water, and blood concentrations in the frontal cortex during sleep within an individual, demonstrating that the greatest changes occur during stage N3 sleep and correlate with EEG slow wave activity. We propose that these observed changes, at least in part, are related to glymphatic activity. Once validated, our approach may have broad implications for the glymphatics field and, more generally, the fields of cerebral fluid dynamics, neuroscience, and neurology.

We observe significantly greater bandpower in the LFO band during N3 sleep for both HbO and water signals, with large effect sizes (Figure 3) compared to the RFO and CFO frequency bands. This is consistent with glymphatic activity, which has been shown to predominantly relate to LFO, or cerebrovascular vasomotor oscillations, generated by the locus coeruleus (Hauglund et al., 2025b). Amplitude quantification in the LFO band demonstrates that water concentrations are elevated in the NREM state compared to wakefulness, confirming the findings reported by Yoon and colleagues (Yoon et al., 2025). Building on this, our analyses demonstrate a sleep stage-specific progression of water concentration, increasing from N2 and peaking in N3. Interestingly, the water concentration positively correlated with delta power (Figure 4B), a relationship significantly enhanced by regressing out the plasma water signal, demonstrating that the coupling between sleep water dynamics and slow-wave activity is strongest for water that is *unrelated* to HbO oscillations. This lends further support to the hypothesis proposed by Yoon and colleagues that their proposed linear minimum mean square error (LMMSE) method for filtering plasma water could result in an improved measure for glymphatics.

Although the most robust signal is observed in the LFO band (Figure 4A), we observed smaller but significant water concentration changes in the RFO band across sleep stages (Figure 6A), again with the greatest concentrations observed in N3 compared to other states. This suggests that pressure dynamics associated with respiration contribute to water concentration changes - our proposed surrogates for glymphatic activity - during deep sleep. Multiple lines of evidence also support that respiratory activity can change intracranial pressure and that breathing can drive CSF oscillations (Vinje et al., 2019; Yildiz et al., 2022). In rodents, this was shown to relate to increased glymphatic activity (Smith et al., 2017). Neither the water concentrations (regressed or not) had a significant relationship with EEG beta. This is contrary to Dagum and colleagues’ findings, where they found beta power was a negative correlate of the impedance reductions hypothesized to represent glymphatic activity (Dagum et al., 2025b). Hauglund and colleagues, however, found no relationship with beta or delta power in their glymphatic clearance measurements. These inconsistencies could potentially be explained by differences in the phenomena being measured, different models (Hauglund used a rodent model), our modest sample size, or the confound of other non-slow-wave-related water contributions, such as intravascular water (plasma) flux.

### 4.2 Water signals are more sensitive than HbO to sleep stage changes and delta power

Across both time and frequency domains, water signals consistently showed stronger modulation across sleep stages compared to HbO. Water signal amplitude, quantified as mean concentration, is elevated during N3 (Figure 4A), again with a robust effect size (\*\*\*\**p* < 0.005, d = 1.31), which could be related to greater expansion in the water compartments, such as the interstitial space, during N3 glymphatic activity. In contrast, HbO mean concentration shows no significant modulation across sleep stages (Figure 4D), with no relationship to delta power (Figure 4E). This asymmetry could be explained by the fact that sleep-related glymphatic activity opens interstitial spaces to significantly increase water concentration, while blood concentration changes are more limited to vascular expansion mechanisms. We took another approach to characterizing water flux by counting significantly elevated peak concentrations (suprathreshold envelope peaks, SEP) instances across the sleep stages (Figure 5), finding these results to be consistent with water concentration analysis. In contrast to LFO and RFO findings, no significant differences were observed in the CFO band (0.6–1.2 Hz) across sleep stages for water or HbO signals. These null findings contrast sharply with the clear stage-specific changes observed in the LFO and RFO. This suggests that water oscillations at the cardiac frequency, although robust in magnitude, do not appear to contribute meaningfully to sleep-dependent modulation of water flux and therefore are unlikely to be major contributors to glymphatic clearance. This is consistent with previous reports of glymphatic activity linking cerebrovascular LFOs and CSF pulsations relating to RFOs.

### 4.5 Implications for monitoring glymphatic function and brain compliance in humans

Using an inexpensive, comfortable headband approach, we observed an increase in water concentration within the frontal cortex correlating with sleep transitions from wake to stage N2 and stage N3 sleep, consistent with glymphatic activity. Further supporting the link between these water measurements and glymphatics is the LFO-predominant effect with some sleep stage contribution in the RFO, as would also be expected given its known impact on CSF oscillations (Nair et al., 2023; Yang et al., 2022). This evidence, combined with the water dynamics related to EEG delta power, offers a compelling case that we are measuring a sleep-active process of water flux in the frontal cortex during deep sleep - most likely glymphatic activity. If this method is validated with invasive gold standard methods (e.g., intrathecal gadolinium with MRI), it could be commercialized, with potential to be employed in large studies characterizing the glymphatic system. This will be critical to establishing the field in the clinical realm. Importantly, the low-cost wearable nature of our approach favors potential for use in the sleep laboratory and at home, offering new insights on sleep physiology. The implications extend beyond sleep health, as glymphatic dysfunction has been implicated in aging (Benveniste et al., 2019; Kress et al., 2014; Yankova et al., 2021), neurodegenerative disease (Ciurea et al., 2023), migraine (Vittorini et al., 2024), stroke (Gędek et al., 2023; Lv et al., 2021), tumor physiology (Ciurea et al., 2023), and brain trauma (Butler et al., 2022; Ferrara et al., 2022; Zhuo et al., 2024). Our approach could be used to benchmark glymphatic activity in these populations, serving as a biomarker for patient stratification, identifying candidates for therapies aimed at enhancing glymphatic activity.

## 5. Limitations and future directions

This study presents a foundational advance in tracking sleep-linked cortical and subarachnoid water fluctuation using a novel water-sensitive fNIRS approach contained in a low-cost wearable headband device, with evidence supporting that this may be a sensitive approach to measuring glymphatic activity and relative changes in brain compliance. However, several limitations warrant consideration. First, our method provides indirect measures of glymphatic activity, relying on measurements of water and blood in the frontal cortex and subarachnoid space. While these metrics are physiologically plausible and align with prior animal and impedance studies, they lack direct validation against the invasive direct measurement using MRI and intrathecal gadolinium. Future work should include cross-modal comparisons to confirm specificity. Second, our use of continuous wave fNIRS inherently has limited specificity. Water and blood measurements are inclusive of the entire photon path, including scalp, bone, subarachnoid space, meninges, blood vessels, and brain. Because the vast majority of the water signal comes from the subarachnoid space, blood vessels, and brain tissue (intracellular and extracellular), differentiating the source of the signal is not possible. We used two strategies to mitigate this: 1) LMMSE regression to filter the water signal related to the blood, subtracting out the hypothesized “plasma water” as described by Yoon and colleagues, and 2) we focused on relative changes across sleep stages where fluctuations are known to occur in both the subarachnoid space and the brain tissue. Importantly, differentiating between changes in the subarachnoid space and cortex is not possible using our methodology. Future work will benefit from adding a short (∼1cm) source-detector signal to regress out scalp-based signal (Saager & Berger, 2005). Third, modeling from other studies using similar continuous wave fNIRS equipment suggests that the photons are unable to penetrate beyond the first centimeter of brain tissue, limiting our ability to measure deeper tissue changes (Strangman et al., 2013b). However, mounting evidence suggests that superficial dynamics may reflect broader cerebral fluid exchange (Dagum et al., 2025a; Iliff et al., 2012; Kiviniemi et al., 2016; Mestre et al., 2018) and our findings show consistent stage-specific signal patterns in these regions. Since glymphatic activity spans both superficial and deep brain compartments, additional studies combining fNIRS with deep-imaging modalities are essential to determine how surface dynamics reflect global fluid exchange. Fourth, our sample size was modest and largely male, limiting generalizability. These results, therefore, require confirmation in larger and more diverse cohorts, particularly those at risk for glymphatic dysfunction, such as individuals with neurodegeneration or sleep disorders. Fifth, our NIRS apparatus was limited in the number of available wavelengths, reducing our ability to measure additional chromophores such as deoxyhemoglobin. For our purposes, focusing on HbO was a reasonable surrogate for “blood” as it is the dominant species, particularly in the more dynamic arterial blood, which enters the brain with greater pressure, displacing water with greater impulse magnitude. Sixth, our use of CW fNIRS does present the risk of total signal attenuation, particularly in the longer 980nm wavelength; however, we ensured each wavelength could be received at the detector with acceptable signal-to-noise ratio measurements at the time of setup (see supplementary material). Seventh, we are unable to verify the photon path taken by our signal, and there is a risk that our measurements originate from non-brain/non-CSF tissue. However, the odds of this are relatively low at the 3-4 cm source-detector distance, and water content is comparatively low in other tissues. Furthermore, our signal nicely correlates with brain activity. Eighth, ∼30–40% of our data was excluded due to motion artifacts, highlighting the need for hardware and algorithmic advancements in future iterations of the system. While strong effect sizes were observed, particularly in low-frequency domains, several comparisons approached but did not exceed traditional significance thresholds. Despite these limitations, the observed patterns support the utility of water-sensitive fNIRS for noninvasive monitoring of sleep-dependent brain fluid dynamics in a naturalistic bedside setting. Our method presents some important advantages over previously reported measurement approaches. Additionally, our signals processing approach includes a detailed characterization of HbO changes, in addition to water, allowing for focused, stage-specific comparisons throughout the course of the night, and it is adaptable to other classification approaches based on EEG or other signals (e.g., sleep depth classifications, heart rate variability, etc). We view this as a foundational step toward optical monitoring of sleep health, with implications in glymphatics, neuroscience, human performance research, neurodegenerative disease, and neurocritical care.

## 5. Conclusion

We demonstrate a noninvasive, inexpensive, real-time optical method for assessing glymphatic-related fluid dynamics during natural sleep in humans. By utilizing a customized water-sensitive fNIRS system and a unique signal processing pipeline, we identified robust, sleep-stage-dependent oscillatory signatures, highly pronounced in SWS, characterized by elevated relative water concentrations, quantified by mean amplitudes, peak frequencies, and peak-to-trough metrics. We propose that these signals serve as proxies for glymphatic activity - largely driven by activity in the LFO spectrum, with some RFO contributions. Future work will include the enhancement of our technology with additional wavelengths and source-detector distances, extending these findings to clinical populations at risk for glymphatic dysfunction, such as individuals with neurodegenerative conditions, trauma, or sleep disorders.

