## Supplementary Document for "Tracking Sleep-Linked Brain Fluid Dynamics Using Modified fNIRS: A Novel Noninvasive Window into Glymphatic Function"

1. **Assessment of Signal-To-Noise (SNR) ratio across the optode array of channels**

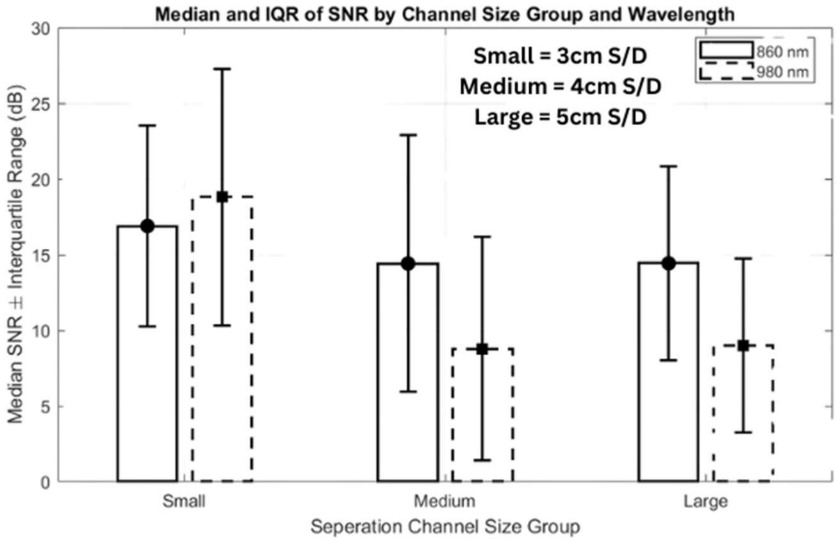


SNR was calculated from the raw optical intensity data during the stable period of each recording. The signal-to-noise ratio (SNR) was computed as the ratio of the power of the signal to the power of the noise, providing a quantitative estimate of data quality across different source–detector (S/D) separations and wavelengths.

As shown in the figure, smaller channel separations (3 cm S/D) exhibited higher median SNR compared to medium (4 cm) and large (5 cm) separations. This trend reflects the expected attenuation of optical intensity with increasing path length through tissue, where longer separations lead to reduced photon detection efficiency and greater susceptibility to physiological and instrumental noise.

Across wavelengths, both 860 nm and 980 nm channels demonstrated comparable SNR, indicating similar overall data quality between the two wavelengths. However, 860 nm channels showed slightly higher median SNR values across all separation groups, likely due to lower scattering and better photon penetration characteristics at this wavelength compared to 980 nm.

2. **Assessment of concentration amplitude changes in LFO (without regressing out the water plasma signal)**

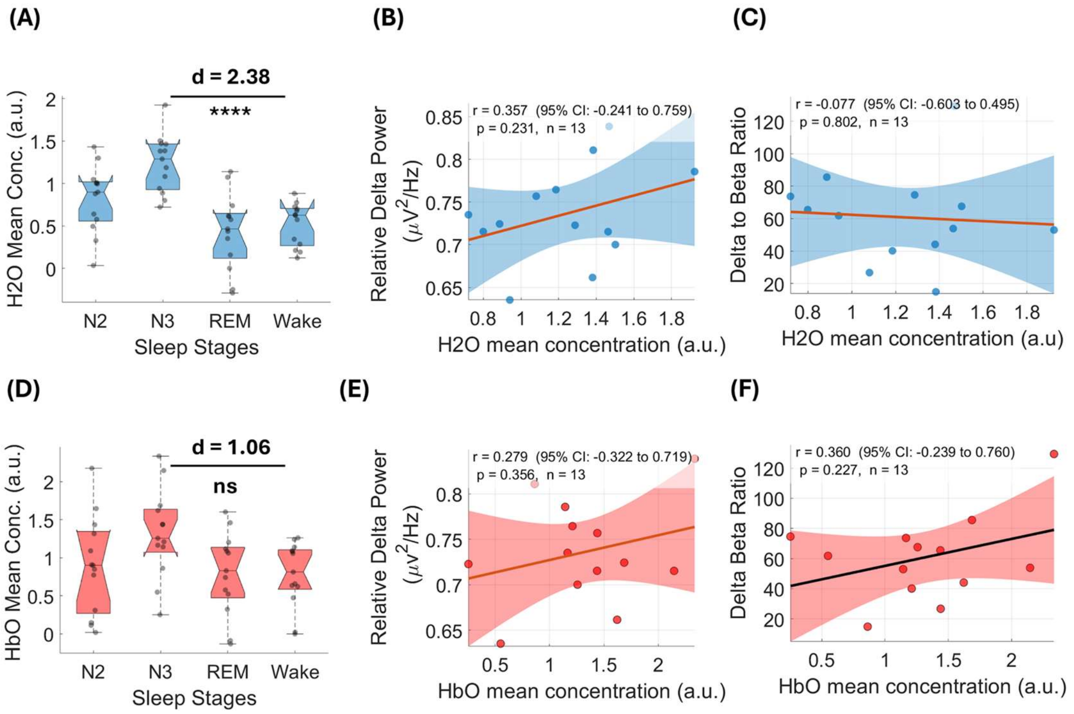


**Supplemental Figure 2. Water and HbO LFO dynamics compared across sleep stages and relationship to EEG power spectra in sleep and wake. (For all plots, N = 9 participants, n = 13 recordings).** (A) H2O relative mean concentration is significantly elevated during N3 relative to other stages. (B) EEG relative delta power may show a trend but does not significantly correlate with H2O mean concentrations. (C) Normalizing EEG delta with beta power demonstrates a slight positive association with H2O relative mean concentrations. (D) HbO mean concentration does not significantly differ across sleep stages or show a significant relationship with EEG delta or delta/beta power (E-F).

The above figure presents an analysis of water and hemodynamic low-frequency oscillations (LFOs) in relation to sleep stages. The figure reveals that the H2O mean concentration is significantly elevated during N3 sleep, though it does not correlate as strongly with EEG markers. The HbO mean concentration and its correlation with EEG markers show no significant changes across sleep stages. These findings suggest that the coupling between brain water dynamics and hemodynamics, rather than just concentration alone, is a sensitive indicator of sleep stage and depth.

3. **Assessment of SEP rate of Water and HbO Signals (without regression of the water plasma signal)**

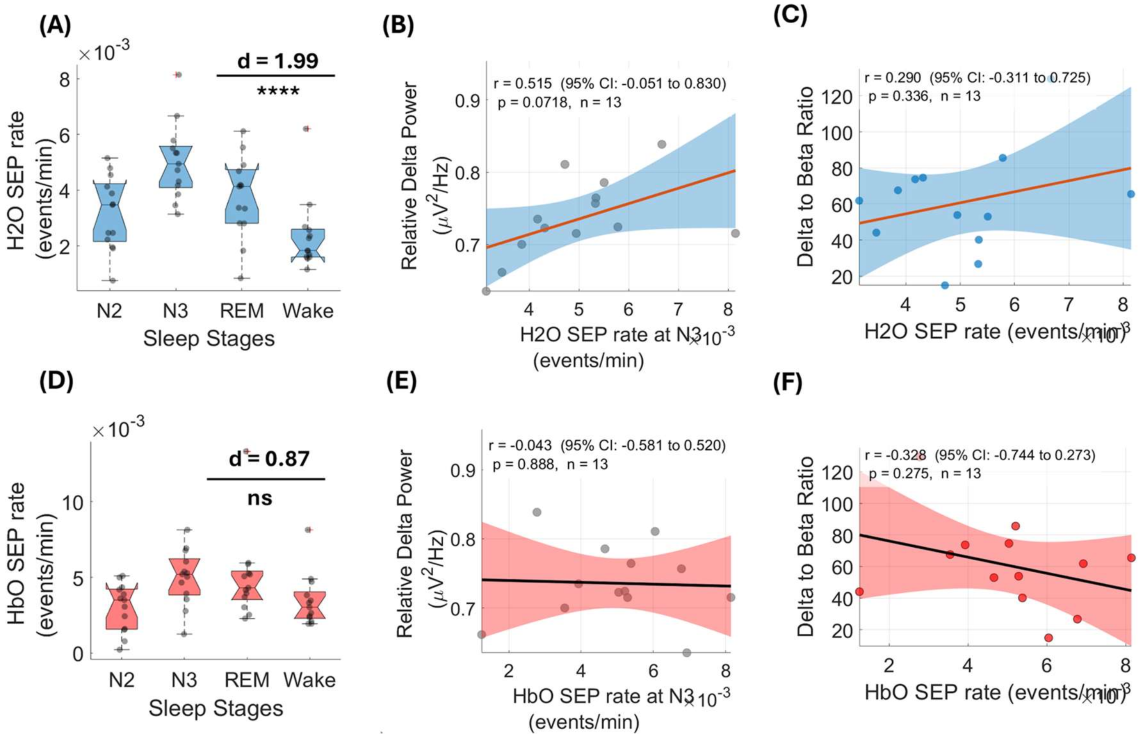


**Supplementary Figure 3: Differential SEP Rates Across Sleep Stages and Correlations with EEG Delta Power.** (A) Box plots showing H2O SEP rates across N2, N3, REM, and Wake stages. (B-C) Correlations between H2O SEP rates during N3 and relative delta power (B) or delta to beta ratio (C). (D) Box plots showing HbO SEP rates across sleep stages. (E-F) Correlations between HbO SEP rates during N3 and relative delta power (E) or delta to beta ratio (F). Significant differences are indicated by asterisks (****p < 0.0001), and effect sizes are shown by Cohen's d.

The figure above illustrates the rates of two different types of Supratheshold envelope peaks (SEPs), H2O and HbO, across various sleep stages. The key finding is a significant increase in H2O SEP rates during N3 sleep (deep sleep), whereas HbO SEP rates show no such stage-specific change. Crucially, these figures were prepared without regressing out the plasma water signal from the total measured water signal. The H2O SEP rates during N3 sleep correlate with markers of deep sleep, such as relative delta power but not so strong correlation with delta to beta ratio. In contrast, HbO SEP rates do not correlate with these markers.

### 4. Assessment of amplitude dynamics in RFO and CFO bands

The figure below compares and mean concentrations across sleep stages for two different regions: RFO (top panel) and CFO (bottom panel).

In the RFO (top row), H2O mean concentration are significantly different across sleep stages, with an increase observed during N2 and N3 sleep. In contrast, the HbO mean concentration shows no significant changes.

In the CFO (bottom row), none of these metrics show a significant difference across sleep stages. The lack of significant changes in H2O mean concentration, and HbO mean concentration in the CFO suggests that the observed sleep-stage-dependent changes are localized to the RFO and not a global phenomenon across the brain. This finding highlights a regional specificity in brain water and hemodynamic dynamics during sleep.

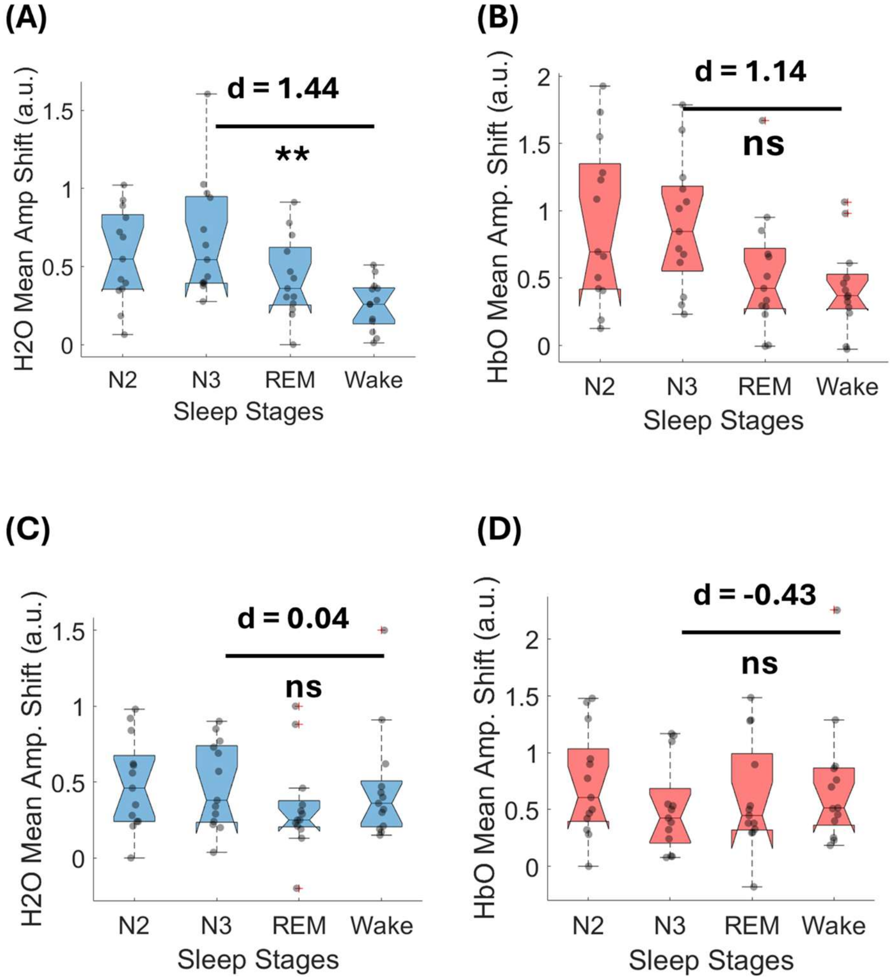


**Supplementary Figure 4: Sleep stage–dependent changes in water and hemoglobin dynamics across RFO and CFO bands.** Panels (A–B) show results in the RFO band, including (A) Mean $H_2O$ concentration, and (B) mean HbO concentration across sleep stages (N2, N3, REM, Wake). While HbO concentration showed a moderate effect (d = 1.14) but did not reach significance, H2O concentration showed significant difference between stage N3 and wake (d=1.44 and p<0.01). Panels (C-D) show results in the CFO band, including (C)Mean $H_2O$ concentration, and (D) mean HbO concentration. No significant differences were observed across sleep stages in CFO. Gray dots represent individual subject values; violin/boxplots depict distributions, median, interquartile range, and 95% confidence intervals.

### 5. Inter-channel variability (without regressing of water plasma signal)

Amplitude analyses revealed clear stage-dependent modulation in water signals when measured at 3 cm source–detector (SD) separation, but much weaker or absent effects at 4 cm separation. Water amplitude shifts were markedly elevated in N3 ($d$ = 2.38, ****,$p$ < 0.0001), consistent with enhanced water dynamics during deep sleep (Panel A). In contrast, HbO amplitude shifts showed no significant differences across stages (Panel C, ns, d= 1.06), highlighting water signals as the more sensitive marker of N3-linked changes. At 4 cm SD, water amplitude shifts were not significant (Panel B, ns), and HbO amplitude effects remained nonsignificant (Panel D, ns). The weaker findings at 4 cm are likely due to a combination of factors. First, roughly 50% of the 4 cm data were rejected during preprocessing, substantially reducing usable trials and statistical power. Second, the device included only four 4 cm channels compared to eight 3 cm channels, further limiting signal averaging and robustness. Third, the signal-to-noise ratio (SNR) of the 4 cm channels was lower than that of the 3 cm channels, which may have obscured subtle amplitude differences. Together, these limitations likely explain why the strong N3-related effects detected at 3 cm were attenuated or absent at 4 cm.

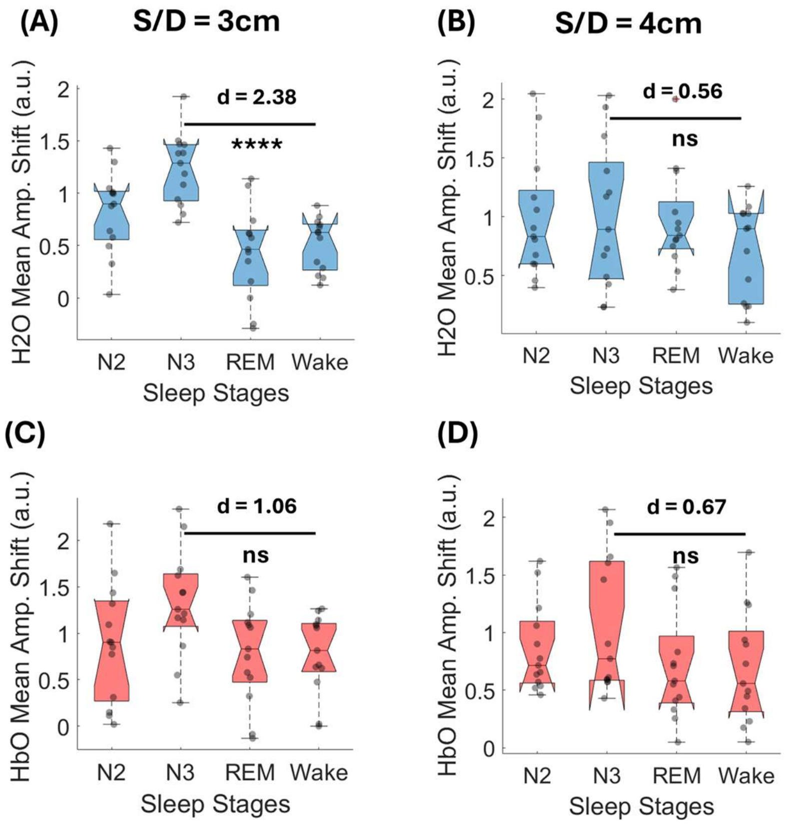


**Supplemental Figure 5: Amplitude shifts in water and HbO signals across sleep stages.**(A) Water ($H_2O$) mean amplitude shifts at 3 cm (B) and 4 cm SD separations. Significant increases were observed during N3 sleep at 3 cm (*d* = 2.38, ****, *p* < 0.0001), but not at 4 cm (*d* = 0.56, ns). (C)) HbO mean amplitude shifts at 3 cm (D) and 4 cm SD separations. No significant differences across sleep stages were found at either distance (*d* = 1.06 and *d* = 0.67, ns).

### 6. Assessment of inter-night variability amongst participants.

To assess stability in NREM-to-Wake phase shift across nights, we quantified both night-to-night variability and coefficient of variation (CV) for each participant. In figure 11, panel A shows residual standard deviation, which reflects the degree of variability across nights. Most participants exhibited relatively low values, suggesting stable measurements, but Participant 11 demonstrated markedly higher variability than all others. Panel B illustrates individual trajectories between Night B and Night C. For most participants, values remained relatively stable across the two nights. However, Participant 11 displayed a sharp increase from Night B to Night C, which drove the elevated variability observed in Panel A. Panel C highlights CV as another index of reliability. Again, Participant 11 stood out with a disproportionately high CV, while the other participants clustered at much lower values, indicating more consistent measurements. These findings suggest that inter-night stability of NREM-to-Wake phase shift is preserved in the majority of participants, but Participant 11 represents an outlier with unusually high variability. This pattern underscores the importance of identifying and accounting for outlier behavior when interpreting individual-level measures of sleep physiology.

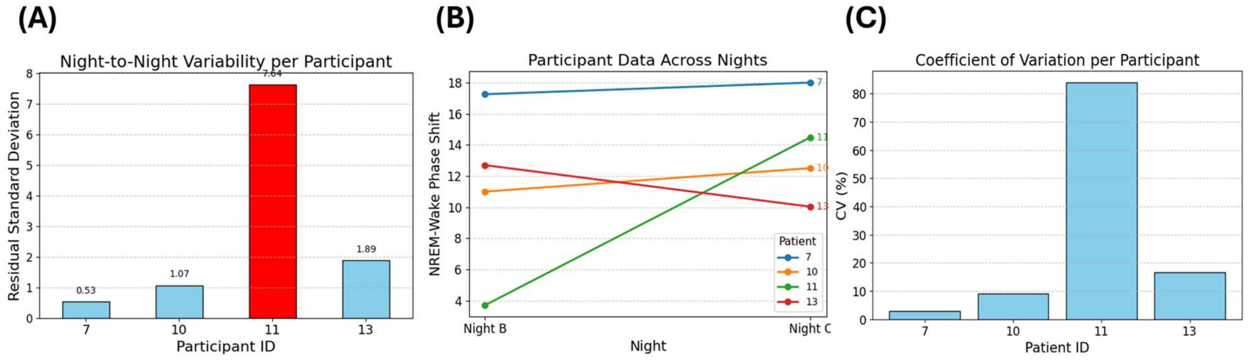


**Supplemental Figure 6: Night-to-night variability and coefficient of variation across participants.** (A) Residual standard deviation illustrating night-to-night variability in NREM-to-Wake phase shift for each participant. Participant 11 showed markedly higher variability compared to others. (B) Individual participant trajectories between Night B and Night C. While most participants showed relatively stable values across nights, Participant 11 demonstrated a large increase from Night B to Night C. (C) Coefficient of variation (CV) per participant, highlighting Participant 11 as having substantially greater variability than others.

### 7. Assessment of Laterality

To assess laterality of the signal across both the hemispheres, we quantified the mean water concentration across the participants corresponding to optodes touching their left and right hemispheres. Analysis of hemispheric water concentration revealed consistent stage-dependent changes across both hemispheres (Supplementary Fig. 7). Relative mean water concentration increased progressively from Wake to N3 sleep, with the highest levels observed during N3 in both left ($d$ = 1.92, $p$ = 5.12 × $10^{-5}$) and right hemispheres ($d$ = 1.90, $p$= = 2.7 × $10^{-5}$). These elevations likely reflect enhanced cortical water accumulation associated with slow-wave activity and glymphatic engagement during deep sleep. The overall pattern and magnitude of change were comparable across hemispheres, suggesting a bilateral process rather than a lateralized effect in sleep-linked fluid dynamics.

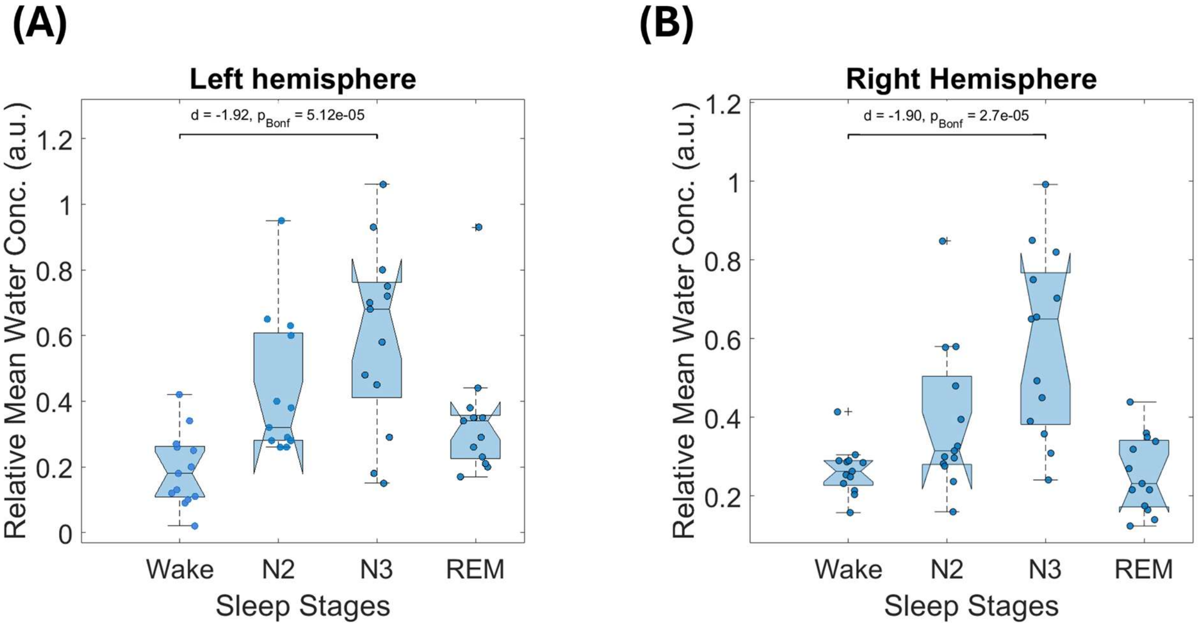


**Supplementary Figure 7: Relative mean water concentration across sleep stages in left and right hemispheres.** (A) Left hemisphere and (B) Right hemisphere show stage-dependent variations in relative mean water concentration (a.u.) across Wake, N2, N3, and REM stages. Both hemispheres exhibit a marked increase during N3 sleep compared to wakefulness (Left: $d$ = 1.92, $p$ = 5.12 × $10^{-5}$; Right: $d$ = 1.90, $p$ = 2.7 × $10^{-5}$), consistent with enhanced cortical water accumulation during slow-wave sleep. Violin plots depict individual data points, median, and distribution spread, illustrating comparable patterns across hemispheres.
