## Supplementary figures and images for "Tracking Sleep-Linked Brain Fluid Dynamics Using Modified fNIRS: A Novel Noninvasive Window into Glymphatic Function"

### Supplementary Figure 1.png

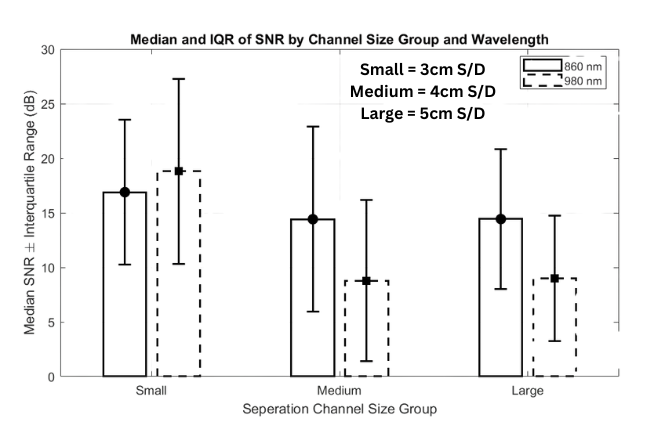

### Supplementary Figure 2.png

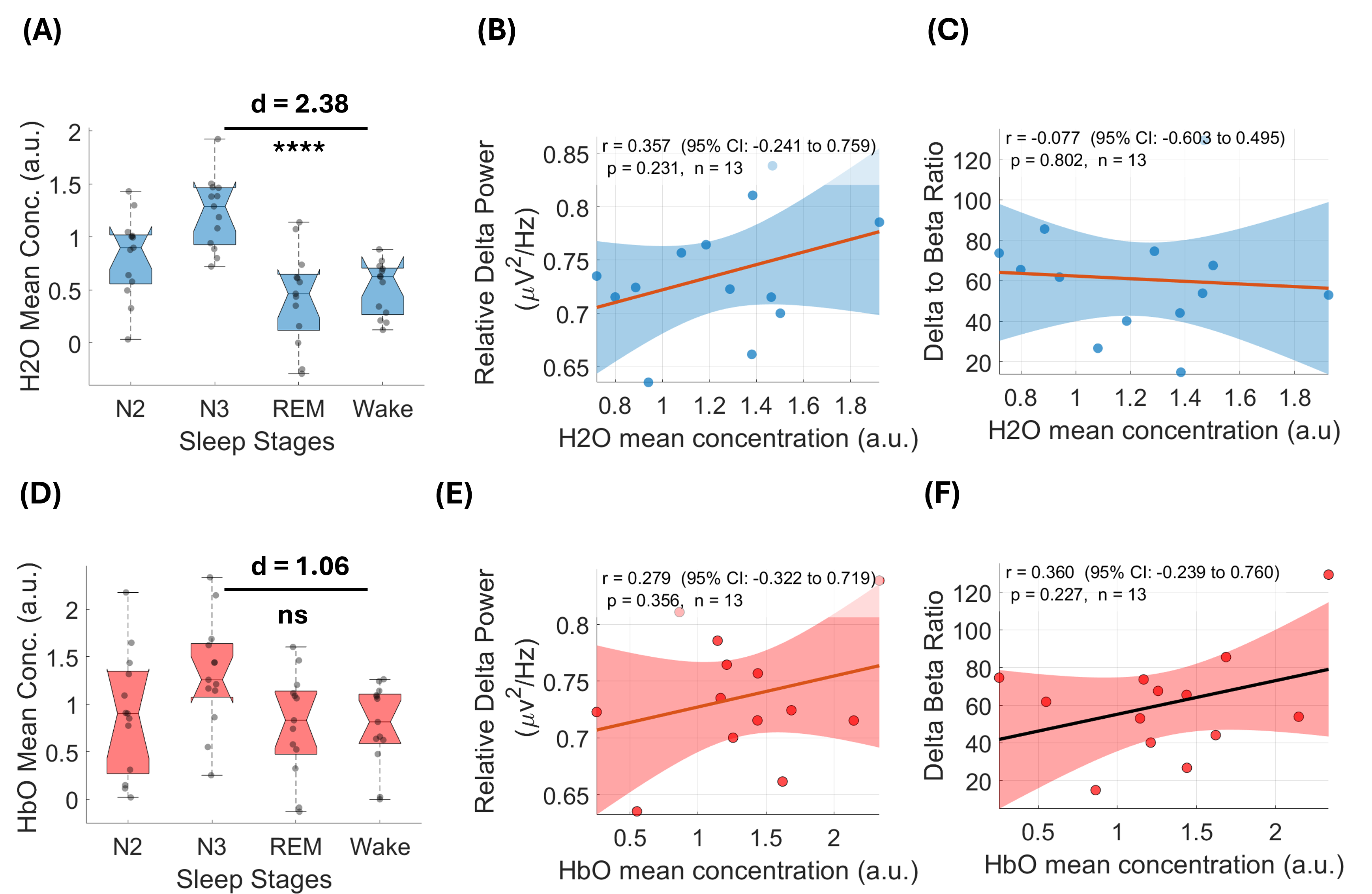

### Supplementary Figure 3.png

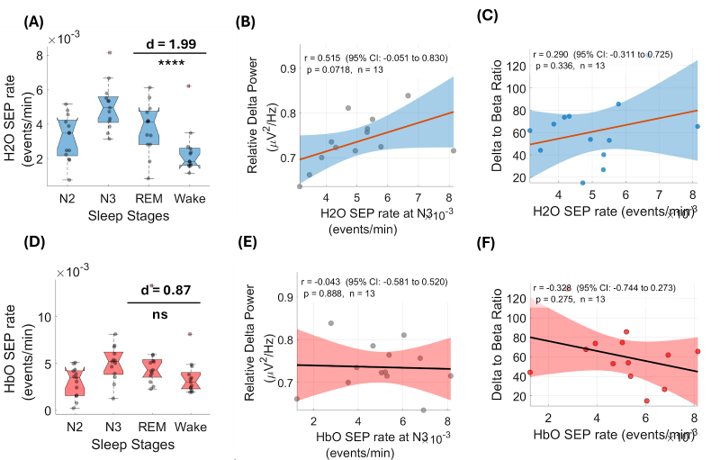

### Supplementary Figure 4.png

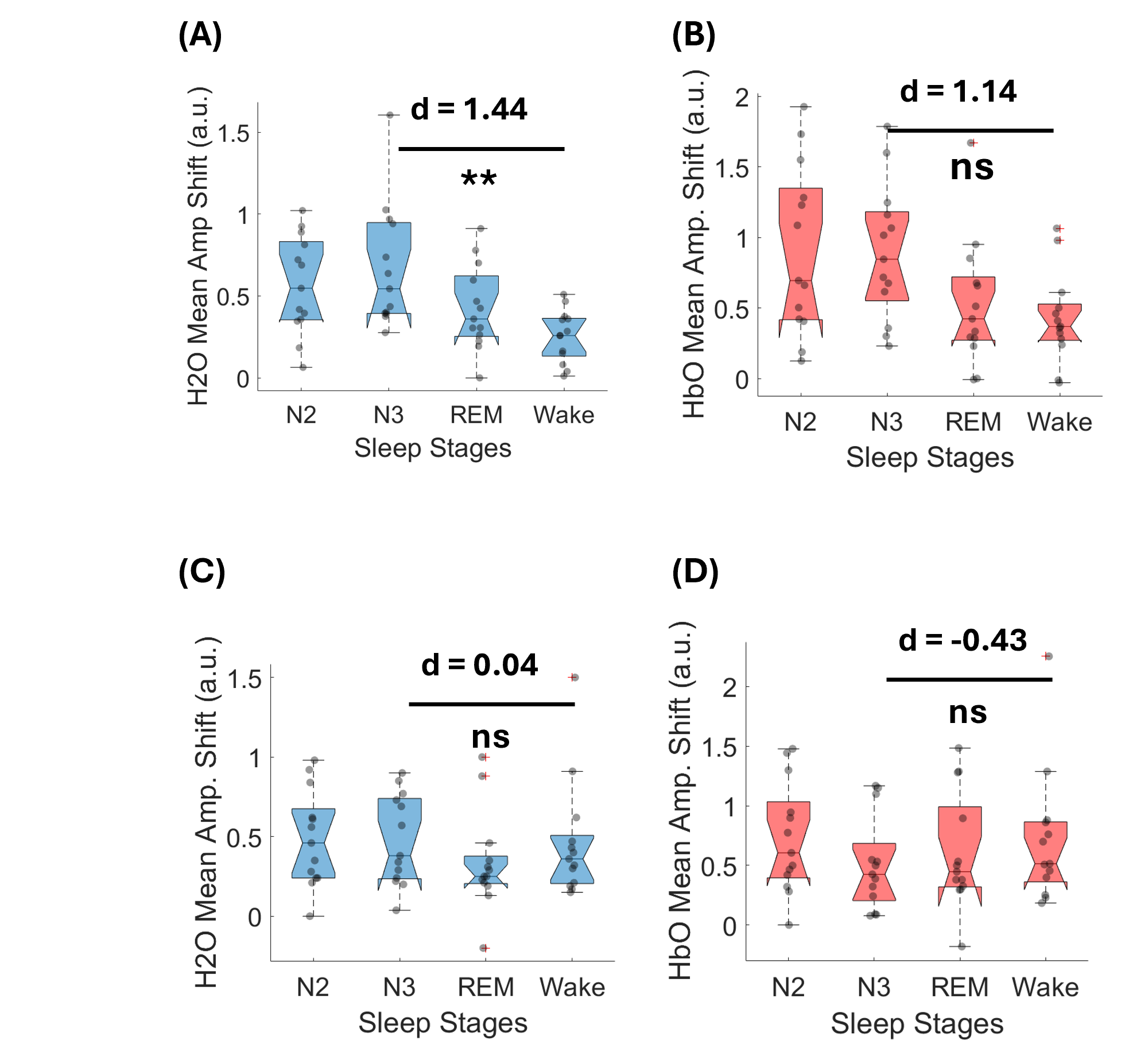

### Supplementary Figure 5.png

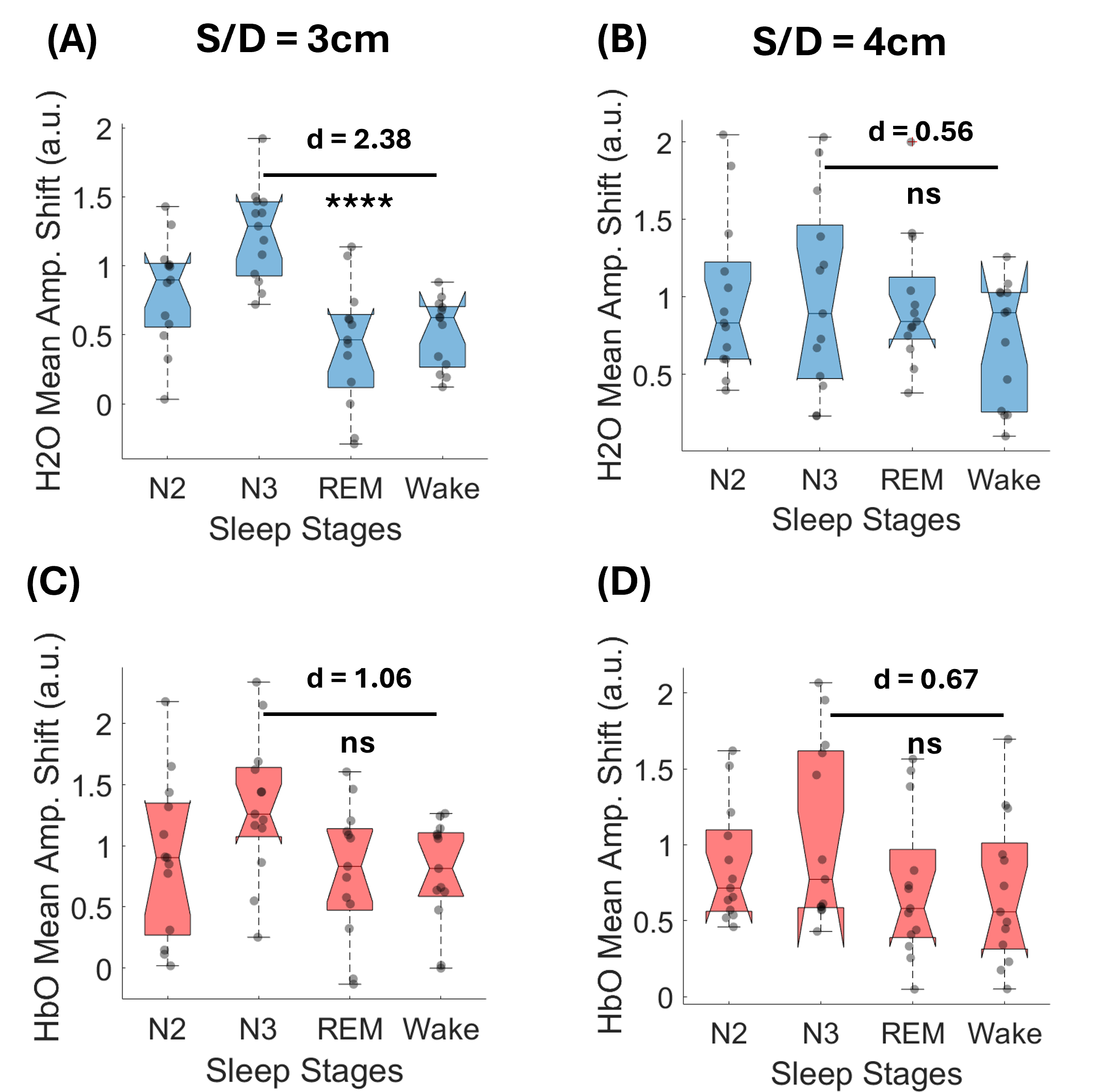

### Supplementary Figure 6.png

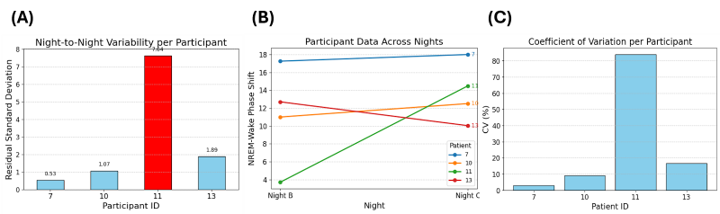

### Supplementary Figure 7.png

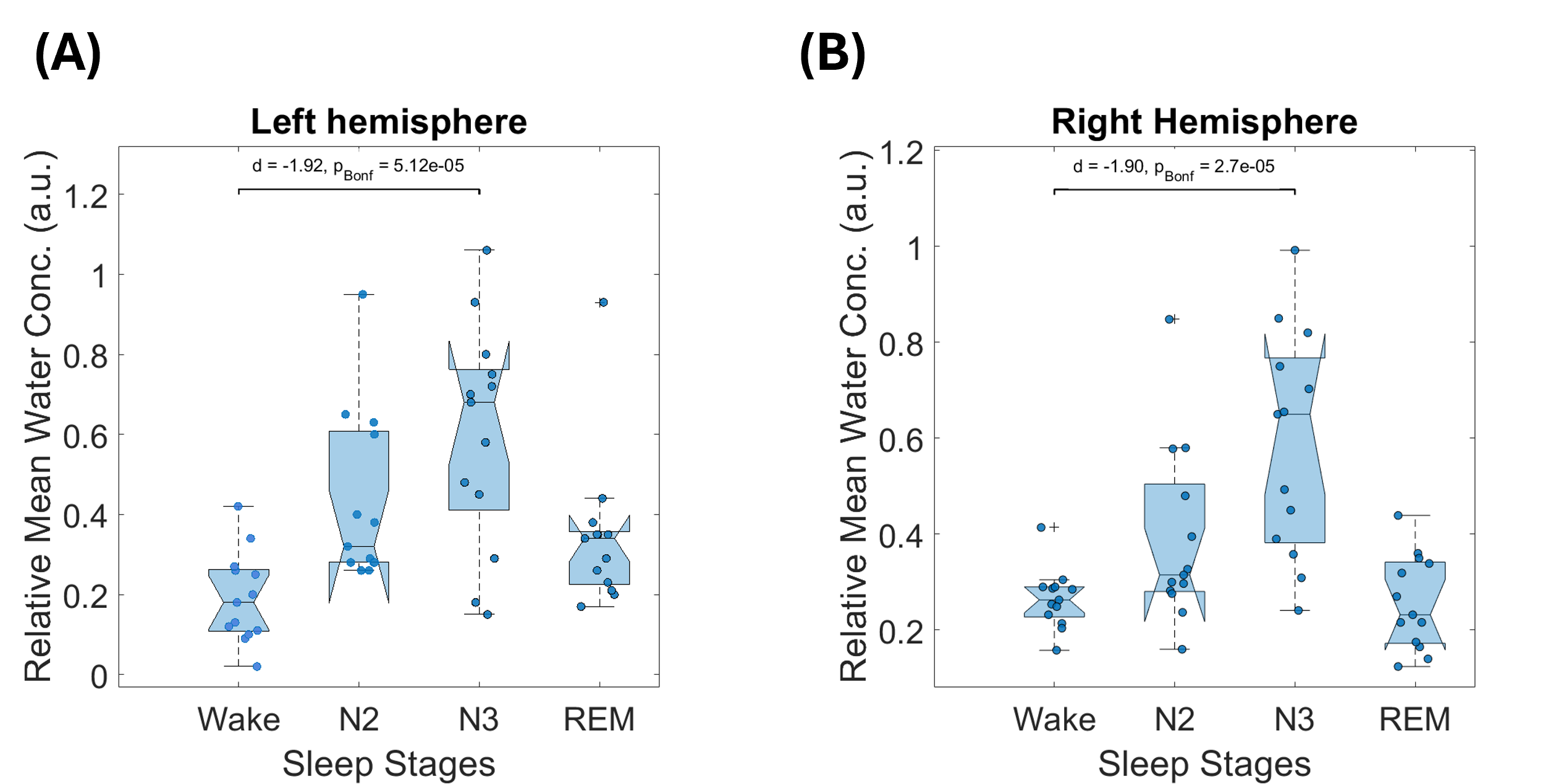
